# Multi-Scale Spiking Network Model of Human Cerebral Cortex

**DOI:** 10.1101/2023.03.23.533968

**Authors:** Jari Pronold, Alexander van Meegen, Hannah Vollenbröker, Renan O. Shimoura, Mario Senden, Claus C. Hilgetag, Rembrandt Bakker, Sacha J. van Albada

## Abstract

Although the structure of cortical networks provides the necessary substrate for their neuronal activity, the structure alone does not suffice to understand it. Leveraging the increasing availability of human data, we developed a multi-scale, spiking network model of human cortex to investigate the relationship between structure and dynamics. In this model, each area in one hemisphere of the Desikan-Killiany parcellation is represented by a 1 mm^2^ column with a layered structure. The model aggregates data across multiple modalities, including electron microscopy, electrophysiology, morphological reconstructions, and DTI, into a coherent framework. It predicts activity on all scales from the single-neuron spiking activity to the area-level functional connectivity. We compared the model activity against human electrophysiological data and human resting-state fMRI data. This comparison reveals that the model can reproduce aspects of both spiking statistics and fMRI correlations if the cortico-cortical connections are sufficiently strong. Furthermore, we show that a single-spike perturbation propagates through the network within a time close to the limit imposed by the delays.

## Introduction

Brain organization and activity display distinct features across multiple spatial and temporal scales: from the molecular level to whole-brain networks, from sub-millisecond processes to memories that last decades (Deco et al., 2008; Honey et al., 2012; Squire et al., 2015). Impressive technological advancements have made almost all these scales accessible through specialized techniques, which leads to a comprehensive but fragmented picture (Sejnowski et al., 2014). Models have the potential to integrate the diverse data modalities into a unified framework and to bridge across the scales (Pulvermüller et al., 2021).

Large-scale, data-driven models at cellular resolution have been constructed for sensory cortex (Reimann et al., 2013; Markram et al., 2015; Girardi-Schappo et al., 2016; Arkhipov et al., 2018; Billeh et al., 2020), prefrontal cortex (Hass et al., 2016), hippocampus (Hendrickson et al., 2012; Bezaire et al., 2016; Ecker et al., 2020), cerebellum (Casali et al., 2019; Yamaura et al., 2020), and the olfactory bulb (Migliore et al., 2014, 2015), among others. These models reproduce resting-state activity (e.g. Potjans and Diesmann, 2014; Markram et al., 2015; Hass et al., 2016; Bezaire et al., 2016) and stimulus responses (e.g. Arkhipov et al., 2018; Billeh et al., 2020) on various levels of detail. Advances in the simulation technology for large networks of point neurons (Einevoll et al., 2019; Jordan et al., 2018; Pronold et al., 2022b,a) have enabled the step beyond single brain regions to multi-area cortical network models (Schmidt et al. 2018a,b; Lu et al. 2022; see also Izhikevich and Edelman 2008 for a pioneering study).

The multi-area spiking network model of Schmidt et al. (2018b) relates the connectivity of the vision-related areas in one hemisphere of macaque cortex to its dynamics. It integrates cortical architecture and connectivity data, in particular axonal tracing data (Bakker et al., 2012; Markov et al., 2014b,a), into a comprehensive, layer-resolved network of 32 areas. Simulations where the model is poised in a metastable regime just below a transition to a high-activity regime reproduce local and cortico-cortical resting-state activity (Schmidt et al., 2018b): single-cell spiking statistics closely match recordings from macaque V1, and functional connectivity patterns correspond well with macaque fMRI data. Moreover, the model yields population bursts that mainly propagate in the feedback direction, akin to empirical findings during visual imagery (Dentico et al., 2014) and in slow-wave sleep (Massimini et al., 2004; Nir et al., 2011; Sheroziya and Timofeev, 2014).

In part due to the scarcity of available human data in comparison with other species, only a few large-scale cellularly resolved human brain network models have been built (Izhikevich and Edelman, 2008; Lu et al., 2022). The former encompasses a million neurons for most simulations (although a variant with 10^11^ neurons was also simulated), while the latter goes up to the full 86 billion neurons of human cortex. The model of Izhikevich and Edelman (2008) includes thalamocortical interactions and displays self-sustained activity as well as chaotic cortical spiking activity (as observed experimentally by London et al., 2010; but see Priesemann et al., 2014). In contrast, Lu et al. (2022) focus on fMRI data and develop a fitting routine to fine-tune the model in order to reproduce recorded BOLD signals. However, both models neglect cytoarchitectural heterogeneity across areas, for instance by using the same average number of incoming synapses per neuron in each brain area. Furthermore, both models simplify laminar patterns of cortico-cortical connectivity, and considerably downscale the number of synapses per neuron. Such downscaling is likely to affect the obtained dynamics, such as the correlation structure of the activity (van Albada et al., 2015).

Leveraging the increasing availability of human data (e.g., Mohan et al., 2015; Minxha et al., 2020; Cano-Astorga et al., 2021; Berg et al., 2021; Shapson-Coe et al., 2021), we build and simulate a model that encompasses the scales from the single-neuron level to the network of areas in one hemisphere of the human brain with a biological number of neurons and synapses in each local circuit. The model aggregates data across many scales, from electron microscopy data for the density of synapses (DeFelipe et al., 2002; Cano-Astorga et al., 2021) to whole-brain DTI and fMRI data, supplements it through predictive connectomics (e.g., Barbas and Rempel-Clower, 1997; Ercsey-Ravasz et al., 2013; Beul et al., 2017; Hilgetag et al., 2019; van Albada et al., 2022), and yields activity data on scales from single-neuron spiking activity to area-level correlation patterns.

Simulating large-scale cellularly resolved models requires the efficient use of supercomputers, a thorough understanding of the inherent bottlenecks of these simulations, and state-of-the-art simulation technology. Systematic benchmarking is a significant step toward the optimal use of neuronal simulator technologies such as NEST (Diesmann and Gewaltig, 2002) on supercomputers (van Albada et al., 2021; Albers et al., 2022). Furthermore, recent studies have systematically isolated and addressed major contributing factors to long simulation times (Pronold et al., 2022b,a). These optimizations, coupled with a relatively coarse cortical parcellation, limit the simulation times for the model presented here. Shorter simulation times lead to a higher turnover rate of simulations, and enable investigations of more versions and realizations of the model.

First, we describe the data integration into a mesoscale connectome, the detailed construction of the model, and the activity data used to validate the model. We validate the mesoscale connectome against features that were not explicitly built in. Then, we analyze the spiking activity in a version of the model with equal local and cortico-cortical synaptic strengths, which we call the ‘ground state’ of the model (note that we do not use the term to refer to an energy minimum). The ground state lacks substantial cortico-cortical interactions, so we systematically increase the cortico-cortical synaptic weights. Next, we compare the resulting activity with single-neuron spiking statistics and area-level correlation patterns based on fMRI; the ‘best-fit’ state is achieved when cortico-cortical synaptic weights are increased relative to local synaptic weights. Finally, we investigate how a single-spike perturbation propagates through the network and find rapid spike propagation in both ground state and best-fit state.

## Materials & Methods

### Model Construction

In the following, we detail the composition of the model and the construction of its ‘mesoconnectome’: the connectivity at the level of neural populations specific to cortical areas and layers. Each of the 34 areas in one hemisphere of the Desikan-Killiany parcellation (Table 1) is modeled as a layer-resolved 1 mm^2^ microcircuit consisting of leaky integrate-and-fire neurons. The layers considered are 2/3, 4, 5, and 6, simplifying laminar subdivisions and ignoring layer 1 in view of its low neuron density. In each local circuit, the full natural density of neurons and synapses for the modeled layers is used. This leads to a total of 3.47 million neurons connected via 42.8 billion model-internal synapses (Fig. 1). The remaining input impinging on the neurons, from non-modeled parts of the brain, is represented as a stochastic drive.

**Table 1.**
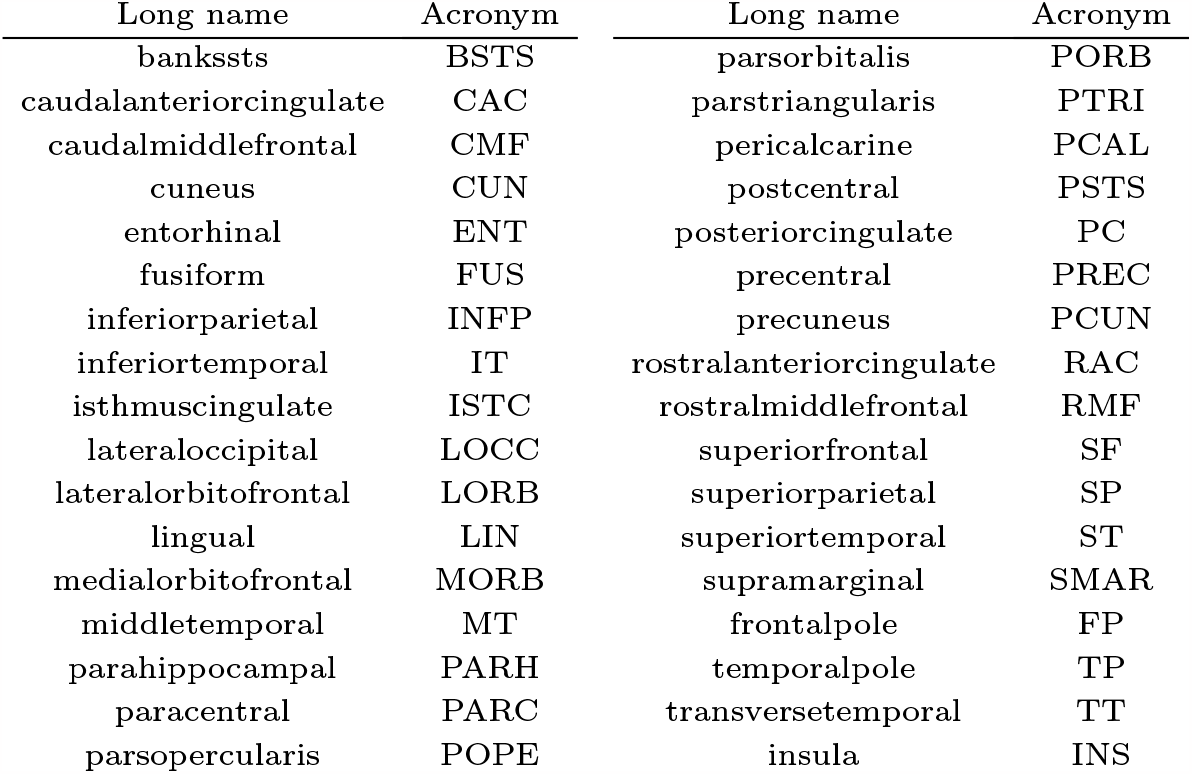
All 34 areas in the Desikan-Killiany parcellation for one hemisphere with corresponding acronyms.

| Long name | Acronym | Long name | Acronym |
| --- | --- | --- | --- |
| bankssts | BSTS | parsorbitalis | PORB |
| caudalanteriorcingulate | CAC | parstriangularis | PTRI |
| caudalmiddlefrontal | CMF | pericalcarine | PCAL |
| cuneus | CUN | postcentral | PSTS |
| entorhinal | ENT | posteriorcingulate | PC |
| fusiform | FUS | precentral | PREC |
| inferiorparietal | INFP | precuneus | PCUN |
| inferiortemporal | IT | rostralanteriorcingulate | RAC |
| isthmuscingulate | ISTC | rostralmiddlefrontal | RMF |
| lateraloccipital | LOCC | superiorfrontal | SF |
| lateralorbitofrontal | LORB | superiorparietal | SP |
| lingual | LIN | superiortemporal | ST |
| medialorbitofrontal | MORB | supramarginal | SMAR |
| middletemporal | MT | frontalpole | FP |
| parahippocampal | PARH | temporalpole | TP |
| paracentral | PARC | transversetemporal | TT |
| parsopercularis | POPE | insula | INS |

**Table 2.**
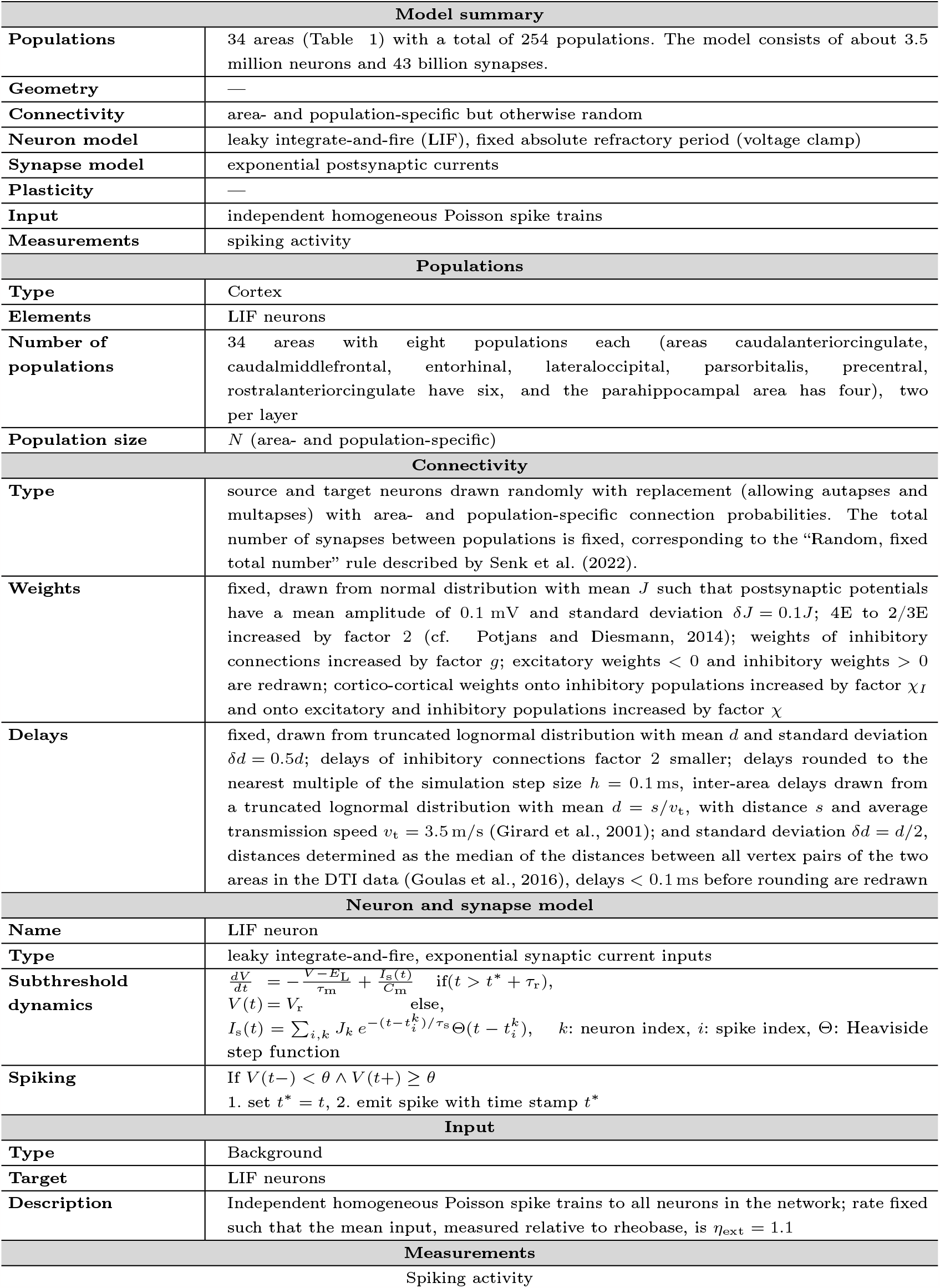
Model description after Nordlie et al. (2009).

| Model summary |  |
| --- | --- |
| **Populations** | 34 areas (Table 1) with a total of 254 populations. The model consists of about 3.5 million neurons and 43 billion synapses. |
| **Geometry** | — |
| **Connectivity** | area- and population-specific but otherwise random |
| **Neuron model** | leaky integrate-and-fire (LIF), fixed absolute refractory period (voltage clamp) |
| **Synapse model** | exponential postsynaptic currents |
| **Plasticity** | — |
| **Input** | independent homogeneous Poisson spike trains |
| **Measurements** | spiking activity |
| Populations |  |
| **Type** | Cortex |
| **Elements** | LIF neurons |
| **Number of populations** | 34 areas with eight populations each (areas caudalanteriorcingulate, caudalmiddlefrontal, entorhinal, lateraloccipital, parsorbitalis, precentral, rostralanteriorcingulate have six, and the parahippocampal area has four), two per layer |
| **Population size** | $N$ (area- and population-specific) |
| Connectivity |  |
| **Type** | source and target neurons drawn randomly with replacement (allowing autapses and multapses) with area- and population-specific connection probabilities. The total number of synapses between populations is fixed, corresponding to the “Random, fixed total number” rule described by Senk et al. (2022). |
| **Weights** | fixed, drawn from normal distribution with mean $J$ such that postsynaptic potentials have a mean amplitude of 0.1 mV and standard deviation $\delta J = 0.1J$; 4E to 2/3E increased by factor 2 (cf. Potjans and Diesmann, 2014); weights of inhibitory connections increased by factor $g$; excitatory weights $< 0$ and inhibitory weights $> 0$ are redrawn; cortico-cortical weights onto inhibitory populations increased by factor $\chi_I$ and onto excitatory and inhibitory populations increased by factor $\chi$ |
| **Delays** | fixed, drawn from truncated lognormal distribution with mean $d$ and standard deviation $\delta d = 0.5d$; delays of inhibitory connections factor 2 smaller; delays rounded to the nearest multiple of the simulation step size $h = 0.1$ ms, inter-area delays drawn from a truncated lognormal distribution with mean $d = s/v_t$, with distance $s$ and average transmission speed $v_t = 3.5$ m/s (Girard et al., 2001); and standard deviation $\delta d = d/2$, distances determined as the median of the distances between all vertex pairs of the two areas in the DTI data (Goulas et al., 2016), delays $< 0.1$ ms before rounding are redrawn |
| Neuron and synapse model |  |
| **Name** | LIF neuron |
| **Type** | leaky integrate-and-fire, exponential synaptic current inputs |
| **Subthreshold dynamics** | $\frac{dV}{dt} = -\frac{V - E_L}{\tau_m} + \frac{I_s(t)}{C_m} \quad \text{if } (t > t^* + \tau_r),$ $V(t) = V_r \quad \text{else,}$ $I_s(t) = \sum_{i,k} J_k e^{-(t-t_i^k)/\tau_s} \Theta(t - t_i^k), \quad k: \text{neuron index, } i: \text{spike index, } \Theta: \text{Heaviside step function}$ |
| **Spiking** | If $V(t-) < \theta \wedge V(t+) \geq \theta$<br>1. set $t^* = t$, 2. emit spike with time stamp $t^*$ |
| Input |  |
| **Type** | Background |
| **Target** | LIF neurons |
| **Description** | Independent homogeneous Poisson spike trains to all neurons in the network; rate fixed such that the mean input, measured relative to rheobase, is $\eta_{\text{ext}} = 1.1$ |
| Measurements |  |
|  | Spiking activity |

**Table 3.** Parameter specification for synapses and neurons.

| Synapse parameters |  |  |
| --- | --- | --- |
| Name | Value | Description |
| $J \pm \delta J$ | Intra-areal connections: $16.4 \pm 1.6$ pA onto excitatory and $7.5 \pm 0.8$ pA onto inhibitory neurons prior to the application of scaling factors,<br>cortico-cortical connections scaled as $J_{cc} = \chi J$ onto excitatory and $J_{cc} = \chi \chi_I J$ onto inhibitory neurons | excitatory synaptic strength |
| $g$ | $g = 5$ | relative inhibitory synaptic strength |
| $d_e \pm \delta d_e$ | $1.5 \pm 0.75$ ms | local excitatory transmission delay |
| $d_i \pm \delta d_i$ | $0.75 \pm 0.375$ ms | local inhibitory transmission delay |
| $d \pm \delta d$ | $d = s/v_t \pm \frac{1}{2} s/v_t$ | inter-area transmission delay, with $s$ the distance between areas |
| $v_t$ | $3.5$ m/s | transmission speed |
| Neuron parameters |  |  |
| Name | Value | Description |
| $\tau_m$ | 10 ms | membrane time constant |
| $\tau_r$ | 2 ms | absolute refractory period |
| $\tau_s$ | 2 ms | postsynaptic current time constant |
| $C_m$ | 220 pF for excitatory neurons,<br>100 pF for inhibitory neurons | membrane capacity |
| $V_r$ | $-70$ mV | reset potential |
| $\theta$ | $-45$ mV | fixed firing threshold |
| $E_L$ | $-70$ mV | leak potential |

**Fig. 1.**
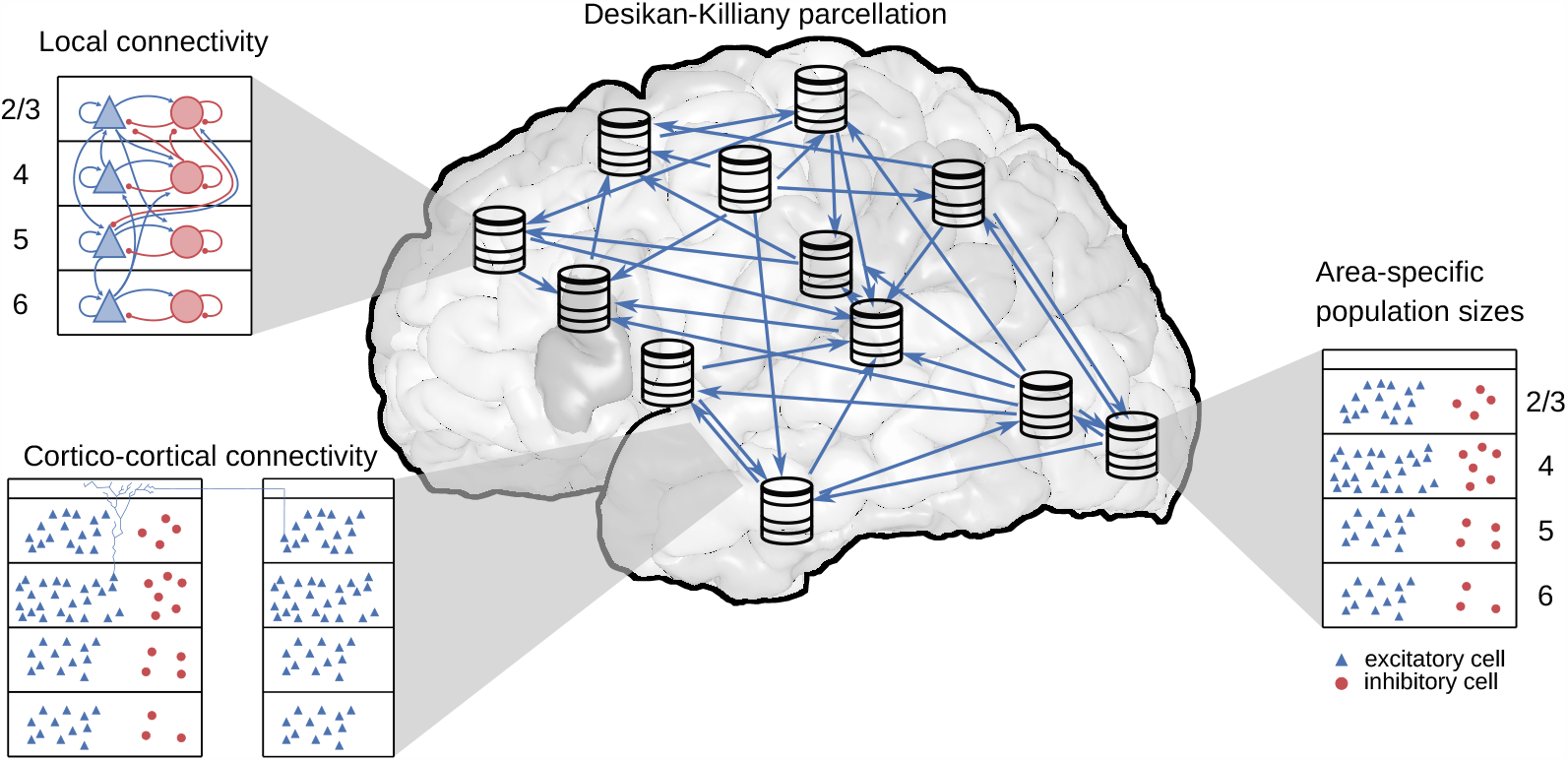
Model overview. The model comprises all 34 areas of the Desikan-Killiany parcellation (Desikan et al., 2006) in one hemisphere of human cerebral cortex. Each area is modeled by a column with 1 mm^2^ cortical surface. Within each column, the full number of neurons and synapses based on anatomical data is included. In total, this leads to 3.47 million neurons and 42.8 billion synapses. Both the intrinsic and the cortico-cortical connectivity are layer- and population-specific.

### Neuron number

The number of neurons per layer follows from multiplying their volume density *ρ*_neuron_ with the layer thickness *h*_layer_ and the surface area *A*_column_ as *N*_neuron_ = *ρ*_neuron_*h*_layer_*A*_column_ (here, *A*_column_ = 1 mm^2^). We use the volume density and the layer thickness provided in the seminal work of von Economo and Koskinas (Von Economo, 2009). This data distinguishes the layers into finer categories than the ones we use in our model. Therefore, we sum the corresponding “layer thickness overall” and average the corresponding “cell content” values weighted by the relative layer thickness.

Furthermore, the data is provided in the parcellation of von Economo and Koskinas; we use the mapping to the Desikan-Killiany parcellation constructed by Goulas et al. (2016, Table 1). In the given mapping, one or more von Economo and Koskinas areas are assigned to each Desikan-Killiany area. For the layer thicknesses, we take the average across the corresponding areas in the parcellation by von Economo and Koskinas (using that the mapping was constructed based on cytoarchitectonic similarity, such that the average is across architectonically similar areas); for the volume densities, we weight the average by the relative thickness of the layers.

To separate the neurons in a given layer into inhibitory and excitatory neurons, we use the layer-resolved relative size of the two populations from the electron-microscopy-based reconstruction of cortical tissue in the human temporal lobe by Shapson-Coe et al. (2021, Supplementary Figure 5B). The resulting fractions of excitatory neurons are 65% in layer 2/3, 79% in layer 4, 78% in layer 5, and 86% in layer 6. The population sizes follow by multiplying the relative population size with the total number of neurons in the layer.

### Synapse number

We approximate the volume density of synapses *ρ*_synapse_ = 6.6*×*10^8^ synapses*/*mm^3^ (Cano-Astorga et al., 2021) as constant across cortex (DeFelipe et al., 2002; Sherwood et al., 2020). This allows us to compute the total number of synapses per area based on their respective volume (Von Economo, 2009), as described above for the number of neurons. The task that remains is to determine the pre- and postsynaptic neurons of these synapses.

### Fraction of Local Connections

We separate the 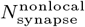 synapses from long-range connections through the white matter from the 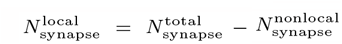 synapses coming from within the area. To determine the fraction of synapses from long-range projections, we use the scaling rule by Herculano-Houzel et al. (2010),

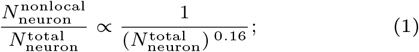

i.e., the relative number of neurons sending axons into the white matter decreases with increasing total number of neurons in the gray matter 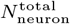. We determine the proportionality constant using the value 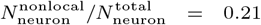 from tracing data in macaque (Markov et al., 2011, though note that this reflects the intra-hemispheric fraction and neglects inter-hemispheric connections) in combination with 1.4 *×* 10^9^ gray matter neurons in macaque (Collins et al., 2010). With the number of gray matter neurons in human, 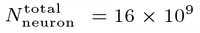 (Herculano-Houzel, 2009), we arrive at the estimate 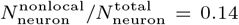 or 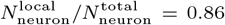. Finally, we assume that the fraction of neurons sending axons into the white matter equals the fraction of synapses from long-range projections, i.e., from cortico-cortical and subcortical sources; in particular, all connections between different cortical areas are treated as white-matter connections.

### Local Connectivity

The 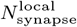 local synapses need a further distinction: 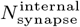 synapses where the presynaptic neuron is part of the simulated column and 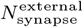 synapses where the presynaptic neuron is outside of the simulated column, i.e., in the remainder of the area. To split these two categories, we use the spatial connection probability *p*(***x***_1_ | ***x***_2_) between a neuron located at ***x***_1_ and another neuron at ***x***_2_, which we assume to be a spatially homogeneous three-dimensional exponential distribution *p*(***x***_1_ | ***x***_2_) *∝* exp(*−*|***x***_1_ *−* ***x***_2_|*/λ*_conn_) with decay constant *λ*_conn_ = 160 *µ*m (Packer and Yuste, 2011; Perin et al., 2011). From *p*(***x***_1_, ***x***_2_) = *p*(***x***_1_ | ***x***_2_)*p*(***x***_2_) where *p*(***x***_2_) is assumed to be constant reflecting a uniform distribution of neurons across space, we obtain the average connection probability *P*_internal_ within the column as

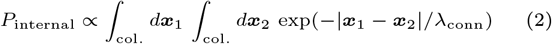

where the proportionality factor is the normalization constant of *p*(***x***_1_, ***x***_2_). We calculate the average connection probability assuming cylindrical columns. In cylindrical coordinates, using *d****x*** = *rdrdϕdz* and 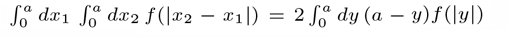 simplifies this integral to

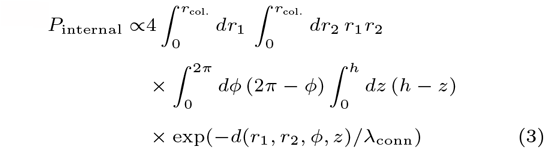

with 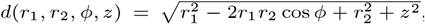, the radius of the column *r*_col._, and the total height of the column *h*. For the probability *P*_external_ that the postsynaptic neuron is in the column but the presynaptic neuron outside of it, the domain outside of the column has to be integrated: ∫_*¬*col._*d****x***_***1***_→∫_*¬*col._*d****x***_***1***_. Approximating the entire area as a cylinder, this leads to the replacement 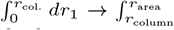 where *r*_area_ is the radius of the larger cylinder, i.e.,

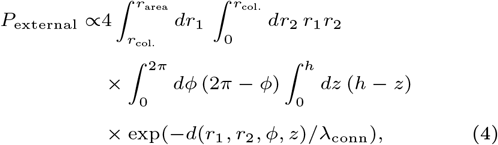

with the same normalization factor as for the internal synapses. The remaining integrals are solved numerically using the adaptive multidimensional quadrature implemented in SciPy (Virtanen et al., 2020). *P*_internal_ and *P*_external_ are used to determine the number of synapses with neurons within and outside of the column, respectively:

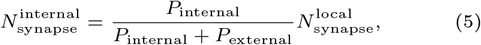

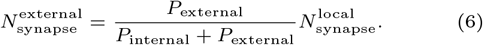

Note that although we keep *r*_col._ the same for all areas, both *P*_internal_ and *P*_external_ are area-specific because their thickness *h*, the total surface area, and the neuron densitiesvary.

For the local connectivity within the column, comprising 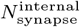 synapses, we use the model of Potjans and Diesmann (2014) as a blueprint. More precisely, we use the average number of synapses 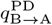 between a neuron in source population B and a neuron in target population A. We combine these average numbers of synapses with the number of neurons 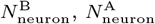 in the pre- and postsynaptic population:

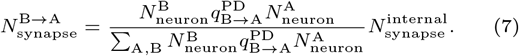

Eq. (7) keeps the relative average number of synapses per pair of neurons (i.e., relative to the other population pairs) equal to the respective value in Potjans and Diesmann (2014) by construction.

The 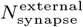 synapses from outside the column are also distributed based on Potjans and Diesmann (2014). Here, we use the indegrees 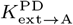 (*k*_ext_(reference) in their Table 5) and the number of neurons in the postsynaptic population 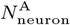 to scale the number of synapses:

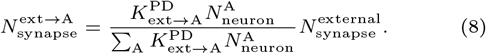

Thus, the external indegrees from Potjans and Diesmann (2014) determine the relative external indegrees for the different populations but not their absolute values. Both in Eq. (7) and Eq. (8), we round the final result to obtain an integer number of synapses.

### Long-range projections

The 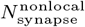 synapses from could belong to intra-or inter-hemispheric cortico-cortical projections, or to projections from subcortical structures. Retrograde tracing in macaque showed that only about 5% of the presynaptic neurons are located in non-adjacent cortical areas within the hemisphere and only about 1% are located in subcortical structures (Markov et al., 2011). Furthermore, contralateral projections (from the other hemisphere) tend to form only a small fraction of the combined cortico-cortical projections (e.g., Dehay et al., 1988; Barbas et al., 2005; Rosen and Halgren, 2022), although this fraction is regionally specific (Ruddy et al., 2017). Based on these observations and the assumption that the fraction of presynaptic neurons equals the fraction of the corresponding synapses, we neglect both subcortical and inter-hemispheric projections, i.e., we treat all 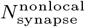 synapses as belonging to intra-hemispheric cortico-cortical projections. Furthermore, we assume that the presynaptic neurons are inside the simulated column in the respective presynaptic area. Thus, we do not consider spatial divergence or convergence of connections beyond the 1 mm^2^ scale.

We define the area-level connectivity according to processed DTI data from Goulas et al. (2016), which is based on data from the Human Connectome Project (Van Essen et al., 2013). For a given target area *X*, we distribute the synapses among the source areas based on the relative number of streamlines NoS_Y*→*X_ in the DTI data,

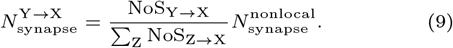

As before, we round the resulting value.

A comprehensive dataset on the layer specificity of the presynaptic neurons based on retrograde tracing is available for macaque (Markov et al., 2014b,a). Not only in this species but also in cat, the layer specificity as measured by the fraction of supragranular labeled neurons SLN, is systematically related to the cytoarchitecture (van Albada et al., 2022). For our human model, we assume the same quantitative relationship as in macaque, for lack of the relevant human-specific data. Fitting a beta-binomial model with a probit link function to the macaque data yields (Schmidt et al., 2018a)

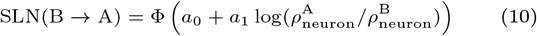

where 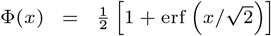 denotes the cumulative distribution function of the standard normal distribution and the fitted parameters are *a*_0_ = *−*0.152 and *a*_1_ = *−*1.534. We use the human neuron densities in Eq. (10) to estimate the laminar origin in human. The SLN value allows determining whether the origin is in layer 2*/*3 or not. Excluding layer 4, which does not form long-range projections (Markov et al., 2014b), the two infragranular layers 5 and 6 still need to be distinguished. To this end, we simply use the relative size of the two populations to distribute the remaining synapses.

On the target side, anterograde tracing can specify the layer specificity. However, there are no comprehensive datasets of anterograde tracing in non-human primates available to date. Hence, we use the collected data from the CoCoMac database (Stephan et al., 2001) which aggregates data across many tracing studies. Relating the target patterns from anterograde tracing to the SLN value reveals three categories of target patterns (Schmidt et al., 2018a):

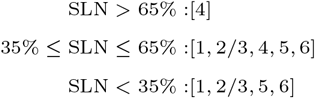

where layer 4 is replaced by 2*/*3 in the first case for agranular target areas (Beul and Hilgetag, 2015). Using the SLN value to distinguish feedforward (SLN *>* 65%), lateral (35% *≤* SLN *≤* 65%), and feedback (SLN *<* 35%) connections, this implies that feedforward connections target layer 4, feedback connections avoid layer 4, and lateral connections show no distinct pattern. For the quantitative distribution of the synapses onto the layers included in the respective target pattern, we use the relative thickness of the layer in relation to all layers of the target pattern.

Thus far, we determined the location of the synapse in the target layer. Next, we decide whether the postsynaptic neuron of a synapse in a given layer is excitatory or inhibitory based on the analysis of the data by Binzegger et al. (2004) in Schmidt et al. (2018a, Table S11). To this end, we sum the target probabilities for postsynaptic neurons across all layers separately for excitatory and inhibitory neurons. This yields the probability for a synapse in a given layer to have an excitatory or inhibitory postsynaptic neuron in any layer. However, we take one exception into account: For feedback connections (SLN *<* 35%), we fix the fraction of excitatory target cells to 93% (Schmidt et al., 2018a) because feedback connections have been found to preferentially target excitatory neurons (Johnson and Burkhalter, 1996; Anderson et al., 2011).

To finally determine the postsynaptic neuron, we assume that all inhibitory postsynaptic neurons are in the same layer as the synapse. For the excitatory neurons, we take the dendritic morphology into account. Using morphological reconstructions of human pyramidal cells in temporal cortex (Mohan et al., 2015) (for a subset of the data see Mohan et al., 2023), we calculate the layer-resolved length of dendrites for neurons with the soma in a given layer. Assuming a constant density of synapses along the dendrites, the ratio of the length *ℓ*_A,B_ of dendrites in layer A *∈* [1, 2*/*3, 4, 5, 6] belonging to neurons with soma in layer B *∈* [2*/*3, 4, 5, 6] to the total length of dendrites in this layer,∑ _B_ *ℓ*_A,B_, determines the probability that the postsynaptic cell is in layer B given that the synapse is in layer A: *P* (soma in B | synapse in A) = *ℓ*_A,B_*/*∑ _B_ *ℓ*_A,B_.

Ultimately, we only need the location of the postsynaptic neuron but not the location of the synapse. Thus, we multiply *P* (soma in B | synapse in A) with the distribution of the synapses across the layers and marginalize the synapse location.

### Further Model Specifications

#### Neuron parameters

We use the leaky integrate-and-fire (LIF) neuron model with exponential postsynaptic currents (Gerstner et al., 2014) for all neurons. To determine the parameter values, we analyzed the LIF models from the Allen Cell Types Database (https://celltypes.brain-map.org/; Teeter et al., 2018; Berg et al., 2021) which were fitted to human neurons. For both excitatory and inhibitory cells, we fix the leak and reset potential to V_L_ = V_reset_ = *−*70 mV. For the threshold potential V_th_, the membrane time constant *τ*_m_, and the membrane capacitance C_m_, we fitted a log-normal distribution using maximum likelihood estimation to the distribution of the respective parameter for all cells in which the LIF model had an explained variance above 0.75 to ensure a good fit of the LIF model. For convenience, we parameterize the log-normal distribution using the mean and the coefficient of variation CV. The resulting mean threshold potential is V_th_ = *−*45 mV for both excitatory and inhibitory cells with CV = 0.21 and CV = 0.22 for excitatory and inhibitory cells, respectively. The resulting mean capacitance is C_m_ = 220 pF and C_m_ = 100 pF with CV = 0.22 and CV = 0.34 for excitatory and inhibitory cells, respectively. To account for the high-conductance state in vivo (Destexhe et al., 2003), we lower the membrane time constant to *τ*_m_ = 10 ms on average with CV = 0.55 and CV = 0.43 for excitatory and inhibitory cells, respectively. We do not distribute the synaptic time constants, which we fix to *τ*_s_ = 2 ms, and the absolute refractory period of t_ref_ = 2 ms.

In all simulations shown in the main text, the neuron parameters are not distributed, i.e., all coefficients of variation were set to CV = 0. Simulations with distributed neuron parameters are shown in the appendix.

#### Synapse parameters

We use static synapses with a transmission probability of 100 %. Excitatory postsynaptic potentials follow a truncated normal distribution with average amplitude 0.1 mV and relative standard deviation of 10 %. The inhibitory postsynaptic potentials also follow a truncated normal distribution with a factor *g* = 5 larger absolute value of the mean and standard deviation. Excitatory (inhibitory) weights are truncated below (above) zero; values outside of the allowed range are redrawn.

Postsynaptic potentials are converted into postsynaptic currents using the conversion factor

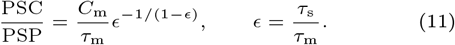

Note that the conversion factor depends on both the synapse parameters (*τ*_s_) and the postsynaptic neuron parameters (*τ*_m_, *C*_m_).

We introduce several scaling factors that affect the postsynaptic potentials: First, the synaptic weights of the synapses within a column from layer 4 excitatory neurons to layer 2/3 excitatory neurons are increased twofold, in agreement with the blueprint (Potjans and Diesmann, 2014). Second, we introduce a scaling factor *χ*_*I*_ for the cortico-cortical synapses targeting inhibitory neurons. This scaling factor stabilizes the column with respect to cortico-cortical input. For all simulations shown in the main text, it is set to 2.0. Third, we introduce a scaling factor *χ* for the cortico-cortical connections onto both excitatory and inhibitory neurons. Increasing this factor leads to the best-fit state reported in Fig. 5 and Fig. 6. For cortico-cortical synapses onto inhibitory neurons, *χ*_*I*_ and *χ* are multiplied with each other.

#### Delays

Within a column, the average transmission delay is 1.5 ms for excitatory and 0.75 ms for inhibitory connections. For the cortico-cortical connections, we assume an average conduction velocity of 3.5 m*/*s (Girard et al., 2001). Dividing the fiber length between two areas, obtained through tractography (Goulas et al., 2016), by this conduction velocity, we obtain the average delay between the two areas. All delays follow a truncated log-normal distribution with a relative standard deviation of 50 %. Delays are truncated below the resolution of the simulation; values outside of the allowed range are redrawn.

#### External input

We determined the number of synapses from non-simulated presynaptic neurons in Eq. (8). The postsynaptic potentials follow a truncated normal distribution with average w_ext_ = 0.1 mV and relative standard deviation of 10 %. We keep the mean input, measured relative to rheobase, fixed at *η*_ext_ = 1.1 and determine the rate of the driving Poisson processes by

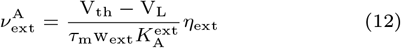

with 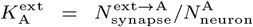. We further introduce two scaling factors for the postsynaptic potentials arriving at excitatory neurons in layer 5 and 6, respectively. For all simulations shown, the first scaling factor is set to 1.05 and the second to 1.15.

### Activity Data

#### Spiking data

Minxha et al. (2020) recorded from thirteen adult epilepsy patients under evaluation for surgical treatment using depth electrodes in medial frontal cortex. In total, they recorded 767 neurons within 320 trials and extracted spikes using a semi-automated spike sorting algorithm. For our analysis, we disregard task-related activity and use only the two seconds of activity which were recorded before stimulus onset. The data is publicly available via the Open Science Framework at http://doi.org/10.17605/OSF.IO/U3KCP.

#### fMRI data

##### Participants

MRI data were obtained from nineteen participants (7 female, age range = 21–33 years, mean age = 25 years) with normal or corrected-to-normal visual acuity. All participants provided written informed consent after receiving full information about experimental procedures and were compensated for participation through either monetary reward or course credit. All procedures were conducted with approval from the local Ethical Committee of the Faculty of Psychology and Neuroscience at Maastricht University. Magnetic resonance imaging

Anatomical and functional images were acquired at Maastricht Brain Imaging Centre (Maastricht University) on a whole-body Magnetom 7T research scanner (Siemens Healthineers, Erlangen, Germany) using a 32-channel head-coil (Nova Medical Inc.; Wilmington, MA, USA). Anatomical data were collected prior to functional data with an MP2RAGE (Marques et al., 2010) imaging sequence [240 slices, matrix = 320 *×* 320, voxel size = 0.65 *×* 0.65 *×* 0.65 mm^3^, first inversion time (TI1) = 900 ms, second inversion time (TI2) = 2750 ms, echo time (TE) = 2.51 ms, repetition time (TR) = 5000 ms, first nominal flip angle = 5^*?*^, and second nominal flip angle = 3^*?*^, GRAPPA = 2]. Functional images were acquired using a gradient-echo echo-planar (Moeller et al., 2010) imaging sequence (84 slices, matrix = 186 *×* 186, voxel size = 1, 6 *×* 1.6 *×* 1.6 mm^3^, TE = 22 ms, TR = 1500 ms, nominal flip angle = 63^*?*^, GRAPPA = 2, multi-band factor = 4). In addition, after the first functional run, we recorded five functional volumes with opposed phase encoding directions to correct for EPI distortions that occur at higher field strengths (Andersson et al., 2003).

Participants underwent five functional runs comprising a resting-state measurement, three individual task measurements, and a task-switching paradigm wherein participants repeatedly performed each of the three tasks. With the exception of the task-switching run, which lasted 9.5 min, all functional runs lasted 15 min. Since task-related runs were not included in the present study, they will not be discussed further. However, it is noteworthy that resting-state runs always preceded task-related runs to prevent carry-over effects (Grigg and Grady, 2010). Participants were instructed to close their eyes during resting-state runs and otherwise to let their mind wander freely.

##### Processing of (f)MRI data

Anatomical images were downsampled to 0.8 *×* 0.8 *×* 0.8 mm^3^ and subsequently automatically processed with the longitudinal stream in FreeSurfer (http://surfer.nmr.mgh.harvard.edu/) including probabilistic atlas-based cortical parcellation according to the Desikan-Killiany (DK) atlas (Desikan et al., 2006). Initial preprocessing of functional data was performed in BrainVoyager 20 (version 20.0; Brain Innovation; Maastricht, The Netherlands) and included slice scan time correction and (rigid body) motion correction wherein all functional runs were aligned to the first volume of the first functional run. EPI distortions were then corrected using the COPE (“Correction based on Opposite Phase Encoding”) plugin of BrainVoyager that implements a method similar to that described in Andersson et al. (2003) and the ‘topup’ tool implemented in FSL (Smith et al., 2004). The pairs of reversed phase encoding images recorded in the beginning of the scanning session were used to estimate the susceptibility-induced off-resonance field and correct the distortions in the remaining functional runs. This was followed by wavelet despiking (Patel and Bullmore, 2016) using the BrainWavelet Toolbox (brainwavelet.org) for MATLAB (2019a, The MathWorks, Natick, MA). Subsequently, high-pass filtering was performed in BrainVoyager with a frequency cutoff of 0.01 Hz and to register functional images to participants’ anatomical images. Using MATLAB, functional data were then cleaned further by regressing out a global noise signal given by the first five principal components of signals observed within the cerebrospinal fluid of the ventricles (Behzadi et al., 2007). Finally, voxels were uniquely assigned to one of 68 cortical regions of interest (ROIs) and an average blood-oxygen-level-dependent (BOLD) signal for each ROI was obtained as the mean of the time-series of its constituent voxels.

### Code & Workflow

The entire workflow of the model, from data preprocessing to simulation and the final analysis, relies on the Python programming language (Python Software Foundation, 2008) version 3.6.5 in combination with NumPy (Harris et al., 2020) version 1.14.3, SciPy (Virtanen et al., 2020) version 1.1.0, pandas (Wes McKinney, 2010) version 0.23.4, Matplotlib (Hunter, 2007) version 2.2.2, networkx version 2.4 (Hagberg et al., 2008), and seaborn (Waskom, 2021) version 0.9.0. All simulations were performed using the NEST simulator (Gewaltig and Diesmann, 2007) version 2.20.2 (Fardet et al., 2021) on the JURECA-DC supercomputer. A simulation of 10 s biological time takes approximately 200 core-hours (1 min build phase + 15 min for 10 s biological time on 768 cores. The workflow is structured using Snakemake (Köster and Rahmann, 2012). For the mean-field analysis, we used the NNMT toolbox (Layer et al., 2022).

## Results

### Human Mesoscale Connectome

The model comprises all 34 areas of one hemisphere of human cortex in the Desikan-Killiany parcellation (Desikan et al., 2006). Each area is modeled by a 1 mm^2^ column and the columns are connected through long-range projections (see Fig. 1). We here give a brief summary of the model construction complementing the details in the Materials & Methods.

We distinguish two classes of neurons, excitatory and inhibitory, and account for the layered structure of cortex. At this level of modeling, the connectivity statistics between neurons in both classes and all layers are needed, which are not straightforwardly delivered by current experimental techniques. Accordingly, we combine available data with predictive connectomics to arrive at a human mesoconnectome at a layer- and population-resolved level. The lack of data on the connectivity is the main reason for considering only two classes of neurons. While a recent study defines 45 inhibitory and 24 excitatory neuron types in human (Hodge et al., 2019), including this diversity would require a huge number of cell-type-specific connection probabilities. This is not yet feasible because no connectivity data is available at such a fine granularity; hence, we restrict the model to two classes of neurons, as done in earlier studies (Potjans and Diesmann, 2014; Schmidt et al., 2018a,b).

#### Mesoscale Connectome

To derive the mesoconnectome, we start from the total number of synapses per layer and subsequently assign pre- and postsynatpic neurons. For the local connections, we use the connection probabilities derived by Potjans and Diesmann (2014) (Fig. 2**A** and Sec. “Local Connectivity”) as a blueprint. The relative connection probabilities across source and target populations are kept constant, and they are only scaled by a constant factor to achieve the desired total number of local synapses in each area. The cortico-cortical connectivity on the area level is specified by DTI data from the Human Connectome Project (Goulas et al., 2016, which is based on the data from Van Essen et al., 2013; Fig. 2**B** and Sec. “Long-range projections”). Synapses associated with long-range projections are assigned to postsynaptic neurons according to morphological reconstructions of human neurons (Mohan et al., 2015; Fig. 2**C** and Sec. “Long-range projections”).

**Fig. 2.**
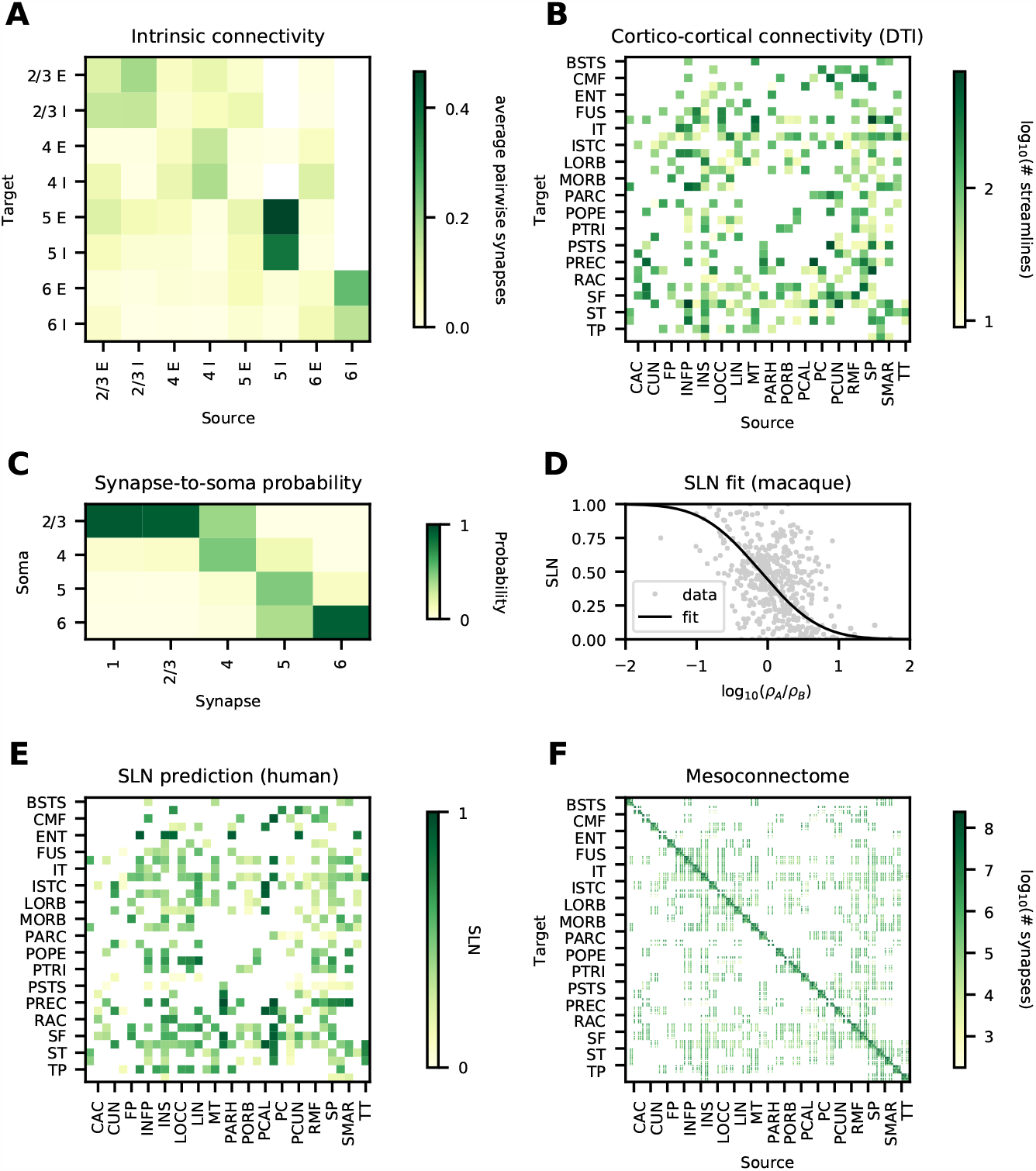
Data and predictive connectomics. (**A**) Within-area connectivity blueprint (average number of synapses per pair of neurons). (**B**) Cortico-cortical connectivity based on DTI (number of streamlines); see Table 1 for acronyms. (**C**) Probability for cortico-cortical synapses in a given layer to be established on neurons with cell body in a given layer, estimated from human neuron morphologies. (**D**) Relation of neuron densities of source area *B* and target area *A* with laminar source pattern (fraction of supragranular labeled neurons, SLN) in macaque. (**E**) Predicted source pattern (SLN) in human. (**F**) Layer- and population-resolved mesoconnectome (number of synapses).

The laminar origin of long-range projections is based on predictive connectomics. Retrograde tracing data in macaque shows that the laminar origin is systematically related to the cytoarchitecture (Hilgetag et al., 2019; Fig. 2**D**). Assuming that the same relation also holds in human, we use the fit in combination with the human cytoarchitecture to determine the laminar origin (Fig. 2**E**). For the laminar target, we assume the same relation between laminar origin and target as done for macaque by Schmidt et al. (2018a), for lack of layer-specific human data.

Combining this data, we arrive at a human mesoconnectome which specifies the number of synapses between excitatory and inhibitory neurons for all areas in the Desikan-Killiany parcellation on a layer- and population-specific level (Fig. 2**F**).

#### Connectivity Validation

To validate the derived mesoconnectome, we compare it with anatomical features that were observed in other species but that were not explicitly built in.

The density of connections between areas is highly heterogeneous, spanning five orders of magnitude, and approximately log-normally distributed in mouse (Gămănut et al., 2018), marmoset (Theodoni et al., 2021), and macaque (Ercsey-Ravasz et al., 2013). Similarly, in our model the numbers of synapses between pairs of populations span five orders of magnitude (Fig. 3**A**) and they are approximately log-normally distributed. Furthermore, the connection density decays exponentially with distance in mouse (Horvát et al., 2016), marmoset (Theodoni et al., 2021), and macaque (Ercsey-Ravasz et al., 2013). In our model, the number of synapses between pairs of areas also decays exponentially (Fig. 3**B**) with a decay constant of 45.6 mm. Thus, two salient features of tracing data are captured by our model.

**Fig. 3.**
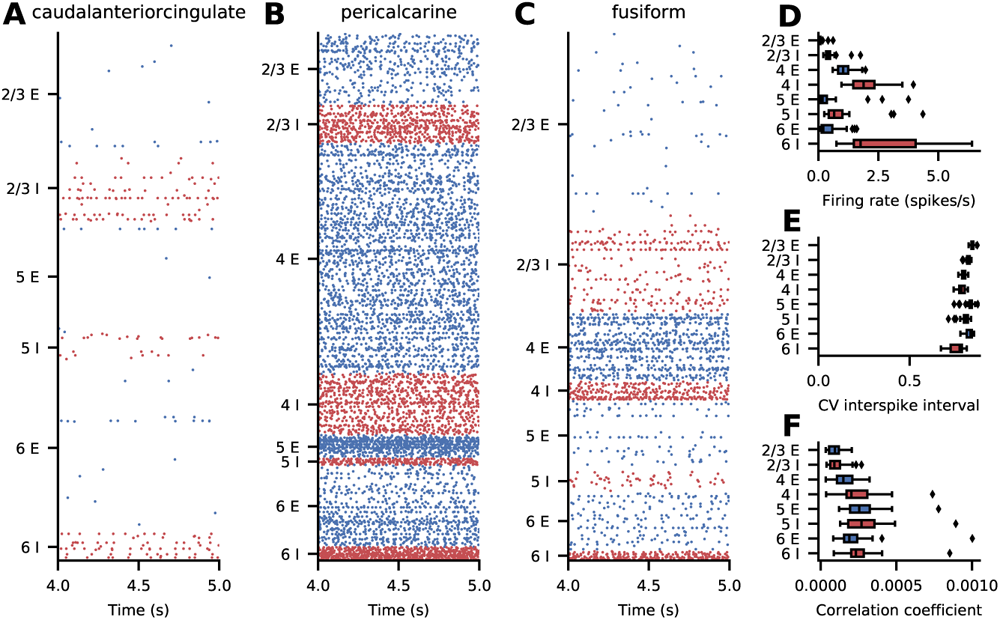
Connectivity validation. (**A**) Histogram of the number of synapses between pairs of populations (gray bars) and a log-normal fit (black line). (**B**) Logarithmic number of synapses between a pair of areas versus distance between these areas (gray symbols) and an exponential fit with decay constant *λ* (black line). (**C**) Average outdegree of a neuron in any given population to any postsynaptic area in either feedforward (FF) or feedback (FB) direction. (**D**) Average number of target areas of a neuron in any given population to any postsynaptic area with average outdegree larger than 100 in either feedforward (FF), lateral (LAT), or feedback (FB) direction.

Anterograde tracing data indicate that feedback axons arborize more strongly than their feedforward counterparts (Rockland, 2019). This suggests a larger outdegree of feedback projections compared to feedforward projections. In our model, the average outdegree from neurons in a given population to a given target area varies systematically between feedforward and feedback projections (Fig. 3**C**); here, feedforward and feedback were classified based on the predicted SLN value (Schmidt et al., 2018a): SLN *>* 65% (feedforward), 35% *≤* SLN *≤* 65% (lateral), and SLN *<* 35% (feedback). The average outdegree for feedforward inter-area connections in our model is 352 compared to 554 in the feedback direction.

Finally, fully reconstructed axons (Winnubst et al., 2019) suggest that many projecting neurons target multiple areas. To check for such divergence in the model, we restrict ourselves to connections with an average outdegree larger than 100. Again using the predicted SLN value to separate feedforward, lateral, and feedback connections, we obtain a broad distribution of the number of target areas (Fig. 3**D**). In addition to the larger outdegree in the feedback direction, feedback projections also target more areas: on average 3.53 compared to 2.46 for lateral and 1.97 for feedforward projections.

### Micro- and Macroscopic Dynamics

#### Ground-state Spiking Activity

We first consider simulations with equal strengths of local and cortico-cortical synapses. The simulated spiking activity of this ‘ground-state’ version of the model is asynchronous and irregular with low firing rates across all areas (Fig. 4). There is a pronounced structure of the activity across populations, layers, and areas (Fig. 4**A**-**C**). To quantify the spiking activity further, we consider population-averaged statistics (Fig. 4**D**-**F**). The firing rate of the inhibitory neurons is higher than the firing rate of the excitatory neurons, with the highest activity in layer 6 (Fig. 4**D**). The activity of some excitatory populations is very low, in particular in layers 2/3 and 5 (Fig. 4**D**). In terms of the irregularity of the spike trains, quantified by the coefficient of variation CV of the interspike intervals, all populations are in the regime of CV ISI *≈* 0.8 (Fig. 4**E**), i.e., slightly more regular than a Poisson process. Lastly, the average pairwise correlation between the neurons is close to zero across all populations (Fig. 4**F**).

**Fig. 4.**
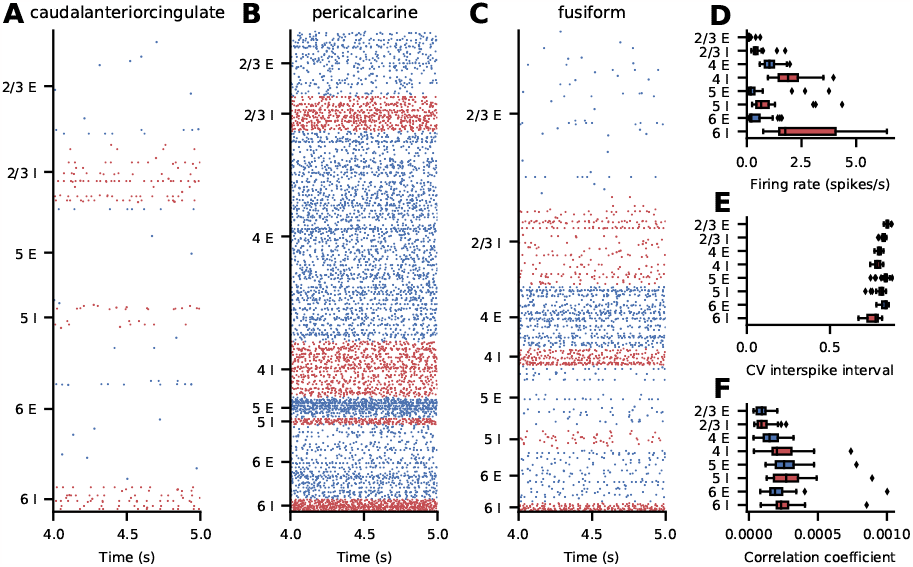
Ground-state spiking activity of the model. (**A–C**) Raster plots for three representative areas; subsampled to 2.5% of the excitatory (blue) and inhibitory (red) neurons. (**D–F**) Layer- and population-resolved distribution of population-averaged statistics across areas; boxes show quartiles, whiskers are within 1.5 times the interquartile range, symbols show outliers outside of the whiskers. (**D**) Firing rate. (**E**) Coefficient of variation of the interspike intervals of neurons with at least 10 spikes. (**F**) Pairwise correlation coefficient of a random subsample of 2000 neurons for each population.

#### Comparison With Experimental Activity Data

To obtain stronger inter-areal interactions, we increase the cortico-cortical synaptic weights onto excitatory neurons by the cortico-cortical scaling factor *χ* and onto inhibitory neurons by a factor *χ*_*I*_ *χ*, where *χ*_*I*_ = 2. We compare the resulting activity of the model with experimental activity data on two levels: on the neuron level, we use the electrophysiological recordings by Minxha et al. (2020) from human medial frontal cortex (cf. Sec. “Spiking data”); on the cortex level, we use resting-state fMRI data from nineteen subjects (cf. Sec. “fMRI data”).

The electrophysiological data were recorded in dorsal anterior cingulate cortex and pre-supplementary motor area; we compare the data with the model activity in area caudalanteriorcingulate. The pre-supplementary motor area overlaps with our model area superiorfrontal but forms only a small part of it, so that the two cannot be meaningfully compared. Since the recordings are not layer-or population-specific, we combine the spike trains of all layers and populations in caudalanteriorcingulate for this analysis. In both the experimental and simulated data, we consider only neurons with at least 0.5 spikes/s for the firing rate, and, for the irregularity, expressed as the coefficient of variation of the interspike intervals (CV) and revised local variation (LvR), we consider only neurons with at least 10 spikes in the respective interval. LvR is a measure of spike train irregularity that corrects for firing rate variations and refractoriness (Shinomoto et al., 2009). As the spike trains comprise only two seconds of activity, we divide the ten seconds of simulated activity into five snippets of equal length. In order to compare the experimental and simulated distributions, we calculate the Kolmogorov-Smirnov distances between them and report 1 *− KS*_dist_ as a measure of similarity, where 0 means no and 1 means perfect similarity. To obtain a proxy for the BOLD signal from our model, we use the absolute value of the area-level synaptic currents (Schmidt et al., 2018b). We compute the functional connectivity using the Pearson correlation coefficient of this BOLD proxy (simulation) or the BOLD signal (experiment). As a measure of the similarity between the modeled and empirical functional connectivity we use the Pearson correlation coefficient and the root-mean-square error (RMSE), in both cases excluding the diagonal where all values are identically one. We convert the RMSE to a similarity measure using exp(*−*RMSE_sim_*/σ*_exp_) where *σ*_exp_ denotes the standard deviation of the functional connectivity. We use both methods because the Pearson correlation is based on relative values and quantifies the linear relationship between the variables, while the RMSE-based measure takes into account the absolute FC strengths.

Fig. 5**A** shows how the different similarity measures depend on the cortico-cortical scaling factor *χ*. The agreements of the CV ISI, the LvR, and the rates initially stay constant and these measures abruptly show a higher agreement at *χ* = 2.5. At *χ*-values close to 2.5, the network sometimes starts in a state of high activity and then, after an initial transient, settles in a lower-activity state or, depending on the random seed, the network operates in a higher-or lower-activity state for the same value of *χ* for the full simulation duration. To exclude transients due to a transition from a high-to a low-activity state, we disregard the first 2500 ms. The distributions of the spiking activity measures continue to match the experimental data well until *χ* = 2.8. Afterwards, the similarities of the irregularity measures CV ISI and LvR deteriorate. The similarity of the fMRI functional connectivity calculated using the Pearson correlation of the experimental and simulated functional connectivity matrices grows from 0.37 to 0.47 and then suddenly drops to 0.33 at *χ* = 2.5, a value around which it remains. The correlation to be maximally accounted for by the model is given by the mean correlation of the experimental functional connectivities across subject pairs, which is 0.63; this ceiling is thus not reached. On the other hand, using the RMSE, the similarity stays initially around 0.38 and grows to 0.46 at *χ* = 2.5, which is consistent with the behavior of the spiking activity measures. The mean RMSE-based similarity between experimental functional connectivities of different subjects is 0.59. Thus, also in terms of this measure, the model does not fully account for the empirical FC structure in human subjects, but it comes closer than the Pearson correlation. As *χ* = 2.5 is the first point at which most measures show good agreement, we use this setting for further analysis. In the following, we refer to this setting as ‘the best-fit state’.

**Fig. 5.**
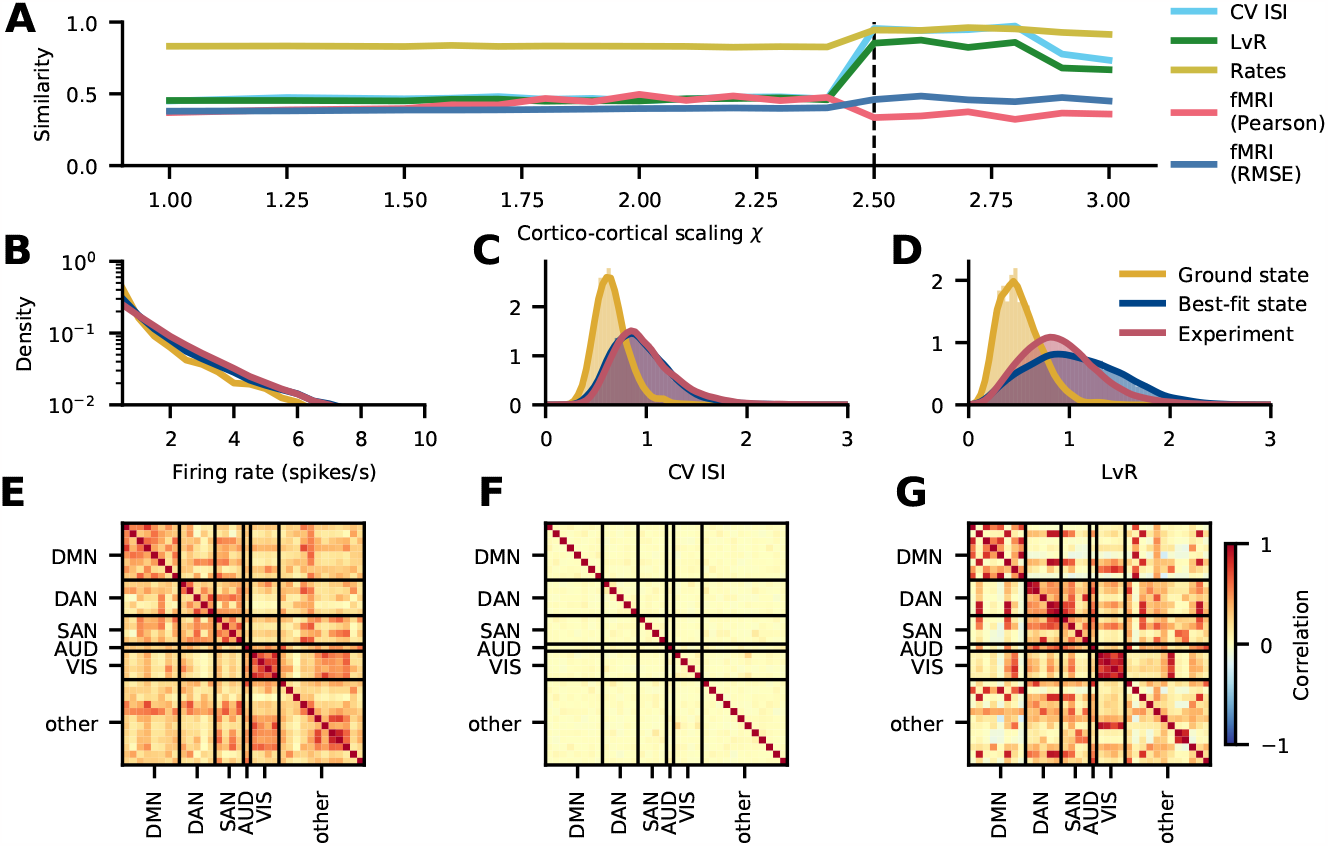
Comparison with experimental activity data. (**A**) Similarity of simulated spiking activity in area caudalanteriorcingulate to experimental spiking data (Minxha et al., 2020) recorded in medial frontal cortex and to resting-state fMRI functional connectivity (cf. Sec. “fMRI data”) as a function of the scaling parameter *χ* for cortico-cortical synaptic strengths. The vertical dashed line at 2.5 corresponds to the chosen best-fit state. (**B–D**) Distribution of spiking statistics across neurons in experimental spiking data (Minxha et al., 2020) and in the simulated ground and best-fit states: Distribution of firing rates (**B**), coefficients of variation of the interspike intervals (CV ISI) (**C**), and revised local variation (LvR; Shinomoto et al., 2009) (**D**). (**E–G**) Functional connectivity in the default mode network (DMN), dorsal attention network (DAN), salience network (SAN), auditory network (AUD), visual network (VIS), and the remaining areas (other). Experimental functional connectivity of the right hemisphere from fMRI recordings, averaged across nineteen subjects (**E**). Simulated functional connectivity based on synaptic input currents in the ground (**F**) and best-fit state (**G**).

A closer look at the underlying statistics (Fig. 5**B-D**) confirms that the best-fit state matches the experimental data better than the ground state does. The firing rate distribution (Fig. 5**B**) is reproduced well by both the ground and the best-fit state, but the latter follows the experimental distribution slightly better. This matches the observation in Fig. 5**A**, where the firing rate similarity is high throughout and peaks at the best-fit state. The CV ISI (Fig. 5**C**) shows clear differences between the ground and best-fit state: in the former, the CV ISI is narrowly distributed around a sub-Poissonian average; in the best-fit state and the recordings, the CV ISI is broadly distributed around a Poissonian average. These two distributions match almost exactly. Similar observations hold true for the LvR, where the main difference compared to the CV ISI is that all distributions are slightly broader.

To facilitate the comparison of the functional connectivities, we group the areas into clusters of different resting-state networks following Kabbara et al. (2017). The experimental (Fig. 5**E**) and best-fit (Fig. 5**G**) functional connectivities show a clear structure with increased correlations within the clusters in the resting-state networks, while the functional connectivity of the ground state shows only very weak correlations (Fig. 5**F**). Also the enhanced correlations between the dorsal attention network (DAN) and the salience network (SAN) are well captured by the model in the best-fit state. These improvements are captured by the RMSE-based measure, which takes into account the absolute FC values, as opposed to the Pearson correlation, which only considers the linear relationship between the empirical and simulated FC.

#### Analysis of Best-Fit State

The simulated spiking activity in the best-fit state varies across areas both quantitatively and qualitatively. Generally, firing rates are higher in the best-fit state than in the ground state (Fig. 6). Some areas, such as caudalanteriorcingulate (Fig. 6**A**) and fusiform (Fig. 6**C**), show low-rate uncorrelated spiking activity with brief population bursts, while some areas, such as pericalcarine, are in a state of high firing in most populations. For completeness, the raster plots of all areas are shown in the Appendix (Fig. 13, 14, 15). We consider population-averaged statistics to quantify the spiking activity on the level of the full network (Fig. 6**D–F**). Inhibitory neurons have higher firing rates than excitatory neurons, with the highest activities in layers IV and VI (Fig. 6**D**). The activity of some excitatory populations is very low, particularly in layers 2/3, 4, and 6. The irregularity of the spike trains, quantified by the coefficient of variation of the interspike intervals (CV ISI), is on average closer to that of a Poisson process compared to the ground state, but also varies more strongly across areas (Fig. 6**E**). The average pairwise correlations are generally low, but reach higher values in a number of areas (Fig. 6**F**).

**Fig. 6.**
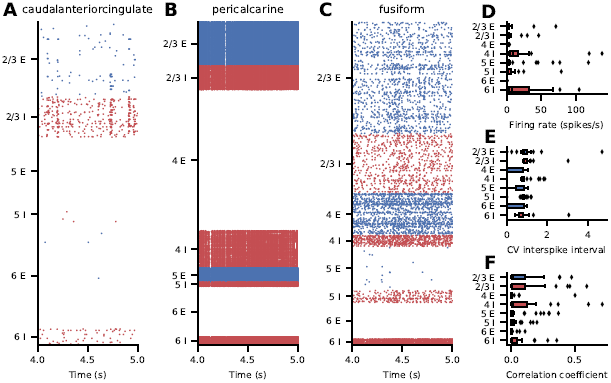
Best-fit spiking activity of the model. (**A–C**) Raster plots for three representative areas; subsampled to 2.5% of the excitatory (blue) and inhibitory (red) neurons. (**D–F**) Layer- and population-resolved distribution of population-averaged statistics across areas; boxes show quartiles, whiskers are within 1.5 times the interquartile range, symbols show outliers outside of the whiskers. (**D**) Firing rate. (**E**) Coefficient of variation of the interspike intervals (CV ISI) of neurons with at least 10 spikes. (**F**) Pairwise correlation coefficient of a random subsample of 2000 neurons for each population.

### Propagation of a Single-Spike Perturbation

In vivo, single-neuron perturbations can affect behavior (Brecht et al., 2004; Houweling and Brecht, 2008). But how does a single-neuron perturbation spread across the cortical network consisting of millions of neurons or more? We investigate this in our model comprising 3.5 million neurons. To this end, we perturb the membrane potential of a single excitatory neuron in layer 4 in primary visual cortex (area pericalcarine) such that it exceeds the threshold and emits a spike. On the network level, this is an extremely weak perturbation. However, since spiking networks are highly sensitive to perturbations (London et al., 2010; Monteforte and Wolf, 2010), even a single spike can alter the spiking pattern of the network (Izhikevich and Edelman, 2008).

We perform two simulations with identical parameters and random seeds but once without and once with the single-neuron perturbation. The drawn random numbers and their total number are the same in both simulations. To quantify alterations of the spiking pattern, we count the total number of spikes of a population in 0.1 ms bins and compute the difference between the unperturbed and the perturbed simulation. As soon as the difference is nonzero due to an additional or missing spike, our observable is set to one. Thus, the observable quantifies the presence or absence of a spike in a given population due to the perturbation. In both the ground state (Fig. 7**A**) and the best-fit state (Fig. 7**B**), the perturbation propagates to all areas in less than 50 ms. In the best-fit state, the perturbation propagates even slighly faster to most areas (Fig. 7**C**). Presumably, the increased activity level in the best-fit state contributes to this difference in propagation speed (Fig. 6). In the ground state, the propagation time is 29.4 *±* 10.9 ms (mean *±* standard deviation); in the best-fit state, it is 25.1 *±* 10.4 ms.

**Fig. 7.**
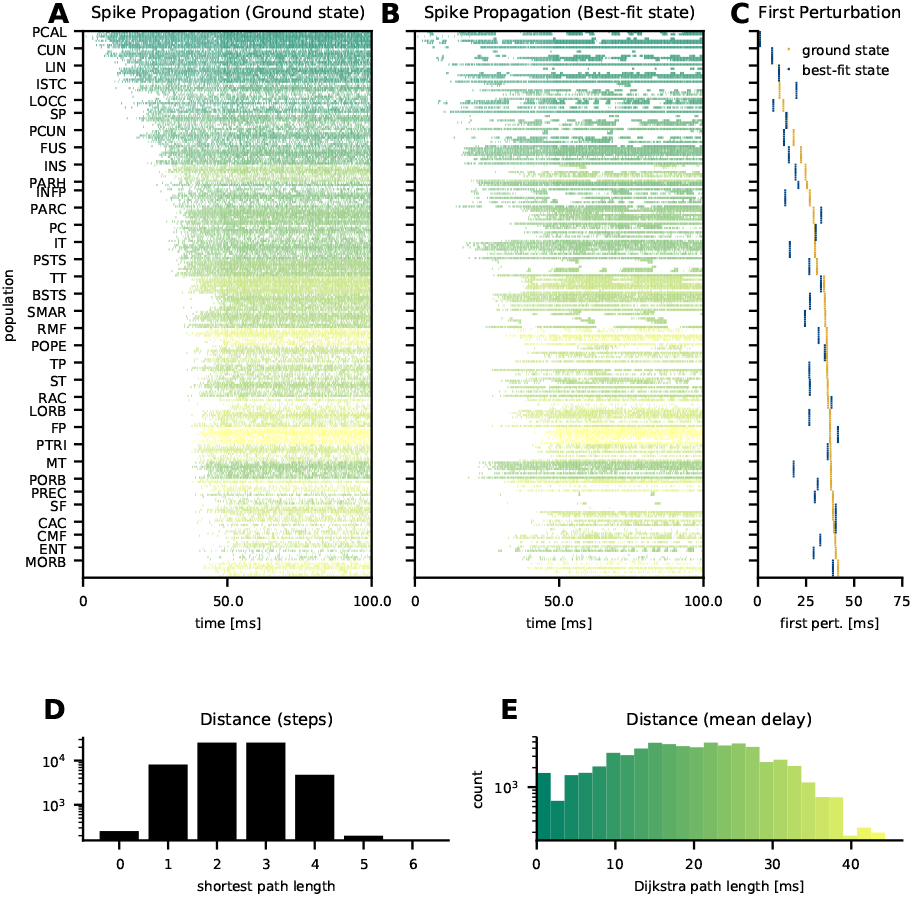
Propagation of the effects of a single spike. Binary absolute difference of spike counts per population in 0.1 ms bins between a perturbed and an unperturbed simulation with identical parameters and random seeds in the ground state (**A**) and the best-fit state (**B**); the color quantifies the Dijkstra path length between the perturbed and the target population. Populations are ordered corresponding to the previous figures; for the scale see panel E. Timing of the first spike count difference per area (**C**) in the ground state (orange) and the best-fit state (blue). Histogram of shortest path lengths between all pairs of populations in the network (**D**). Histogram of shortest path lengths weighted by the average delay between all pairs of populations in the network (**E**).

How is this fast propagation possible? Just like weighted area-level cortical graphs of mice and macaques (Bassett and Bullmore, 2017), the population-level graph in our model exhibits small-world network properties (Watts and Strogatz, 1998). Namely, only a small number of steps is needed to reach any node: the shortest path length between any pair of populations is between one and four and at most five (Fig. 7**D**). But the shortest path length in terms of the number of populations traversed does not account for the transmission delay, which is particularly relevant between areas. Taking also the delay into account by weighting each step with the mean delay and computing the Dijkstra path length (Dijkstra, 1959), i.e., the shortest path based on the sum of the mean delays, we see that the small-world property of the network enables a Dijkstra path length below 50 ms for any pair of populations and below 40 ms for the majority of pairs (Fig. 7**E**). Thus, the network structure supports fast propagation at the population level. The propagation of the perturbation indeed takes place on a timescale similar to the Dijkstra path length between the perturbed population and the target population. The distribution of delays (present in both model versions) in principle allows propagation to take place even faster than this path length.

## Discussion

We aggregated data across multiple modalities, including electron microscopy, electrophysiology, morphological neuron reconstructions, and diffusion tensor imaging (DTI), to construct a multi-scale spiking network model of human cortex. In this computational model featuring 3.5 million neurons connected via 43 billion synapses, each area in a full hemisphere of human cortex is represented by a millimeter-scale layer-resolved microcircuit with the full density of neurons and synapses. The model was simulated on a supercomputer, making use of advances in the simulation technology of NEST. We filled gaps in the data using statistical regularities found in other species, in particular to determine the laminar origins and targets of cortico-cortical connections. Comparisons with electrophysiological recordings from human medial frontal cortex and human fMRI reveal that the model captures aspects of both microscopic and macroscopic resting-state activity when the strength of the cortico-cortical synapses is increased.

Simulations of the model with equal local and cortico-cortical synaptic strengths reveal a state with asynchronous and irregular activity, which we refer to as the ‘ground state’ of the model. The activity is heterogeneous across areas, layers, and excitatory and inhibitory populations. The ground-state activity deviates from the experimental recordings in terms of both spiking activity and inter-area functional connectivity. On the single-neuron level, the distribution of the spiking irregularity in the model is more narrow than the observed one and centered in the sub-Poissonian regime. On the network level, the activity is hardly correlated between areas, which stands in stark contrast to the salient structure in the fMRI data.

To alleviate these discrepancies, we increased the synaptic weights of cortico-cortical connections. The increased anatomical connection strength leads to an increase in inter-areal correlations, with a modular structure similar to the experimental data. On the level of the single-neuron statistics, the increased cortico-cortical synaptic weights hardly affect the distribution of firing rates and irregularity until the synaptic weights reach a critical value at which the fit to the experimental data suddenly improves. This best-fit state features not only stronger correlations between the activity in different areas but also within areas and populations. Furthermore, the firing rates, in particular in the inhibitory populations, are increased. Although the low overall firing rates and the higher inhibitory compared to excitatory rates are realistic features (Dąbrowska et al., 2021; Dehghani et al., 2016), some layers and populations of the model exhibit either excessive or nearly vanishing rates. This uneven distribution of spiking activity across layers and populations remains to be addressed in future.

Computational models allow one to examine questions that are hard to investigate experimentally. Here, we use our model to track the effect of a single additional spike through the large-scale network. We find that the single-spike perturbation spreads across the entire network within less than 50 ms, close to the lower limit imposed by the mean transmission delay between the areas along the shortest possible path. In the best-fit state the propagation is even faster than in the ground state. The observed latencies are on the same order as visual response latencies across macaque cortex (Lamme and Roelfsema, 2000), but note that single-spike perturbations may not be visible on the population level. Rapid propagation of spiking activity, whether on the single-neuron or the population level, is likely to support fast sensory processing and behavioral responses. Due to the stochastic input to the network and its sensitivity to small perturbations, the triggered spike sequences are not fixed but will differ between trials. Future work may investigate whether subnetworks with strong synapses, such as those modeled for turtle cortex by Riquelme et al. (2023), can support repeatable and precisely timed spike sequence in the human cortical network.

The approach we followed closely resembles that taken for the multi-area model of macaque vision-related cortex of Schmidt et al. (2018a,b). A notable difference compared to that model is that our best-fit state is stable over the full length of the investigated simulations, in contrast to the metastable activity obtained there, which sometimes switched to a high-activity state after long simulation durations. In the best-fit state, our model still exhibits a type of metastability: in some simulations, the activity is initially high and later switches to the lower-activity state that matches the experimental data better and that we analyze. The increased stability of the best-fit state in the present model compared to the macaque model and the lack of excessive network-averaged firing rates throughout the simulations provide a better match to actual brain activity.

Just like the model of Schmidt et al. (2018a,b), the present model predicts that stronger cortico-cortical compared to local synapses are needed to account for appreciable functional connectivity between areas, a feature that may be investigated experimentally. In our model, the cortico-cortical synapses are, moreover, stronger onto inhibitory than onto excitatory neurons. A similar feature was reported in mice, where interareal excitatory synaptic input to layer 2/3, but not to layer 5, parvalbumin-expressing interneurons is stronger than to pyramidal neurons (D’Souza et al., 2016; Yang et al., 2013; D’Souza and Burkhalter, 2017). However, using estimates of the relative densities of excitatory and inhibitory neurons taken from cat area 17 (Gabbott and Somogyi, 1986; Binzegger et al., 2004; Potjans and Diesmann, 2014), we were also able to obtain good correspondence with experimental resting-state activity in simulations with very strong cortico-cortical synapses, equal in strength onto excitatory and inhibitory neurons (Fig. 12). In this case, stability was afforded by stronger local synapses onto inhibitory compared to excitatory cells, consistent with slice data from human cortex (Campagnola et al., 2022). In all cases, we did not need to adjust the connection densities to obtain plausible activity as done in the macaque model (Schuecker et al., 2017). This is an improvement because now the connection densities can be directly estimated from the empirical data.

A question that naturally emerges is what sets human cortex apart from that of other species in terms of the properties that determine its resting-state activity statistics. One property that differs with respect to other species is the fraction of excitatory vs. inhibitory neurons, which appears to be lower especially in human cortical layer 2/3 (Gabbott and Somogyi, 1986; Sahara et al., 2012; Shapson-Coe et al., 2021; Alreja et al., 2022). Our model predicts that this reduced excitation in the supragranular layers necessitates greater inter-area coupling for the resting-state activity statistics to match the experimental data, and further leads to a slighty different pattern of functional connectivity between areas (cf. Fig. 5, Fig. 12). Future work may consider a selective increase in the occurrence of bipolar-type interneurons, which preferentially target other inhibitory neurons (Loomba et al., 2022). Further, human cortical neurons tend to be larger and have a lower count density than in other species, receiving more synapses per neuron on average (Sherwood et al., 2020; Loomba et al., 2022). This is likely to be advantageous for information processing, due to a combinatorial explosion of potential synaptic co-activations, but even the implications for resting-state activity remain to be understood. As we have also mentioned and incorporated into our model, the inter-area connectivity of human cortex is sparser because the increased surface between the gray and white matter does not make up for the increased brain volume, so that relatively fewer myelinated axons can connect the areas than in species with smaller brains (Herculano-Houzel, 2009). Another prominent feature of human cortex is its large number of areas, although the increase in this number with respect to other species appears only moderate compared to the expansion of the surface area (Changeux et al., 2021). The current study uses a coarse parcellation both for computational efficiency and to limit the number of unknown parameters, but future work may refine the model toward the potentially 180 or more areas per human cortical hemisphere (Glasser et al., 2016; Amunts et al., 2020). A further aspect, not yet considered here, is the large transcriptional diversity of human cortical neurons, which putatively form hundreds of cell types (Hodge et al., 2019; Miller et al., 2019). Taking into account this extensive diversity would necessitate estimating a huge number of connection probabilities, scaling with the square of the number of cell types, which the available experimental data do not yet allow. This complexity may be gradually approached in future. Also certain electrophysiological properties differ between the cortical neurons of humans and those of other species; for instance, human layer 2/3 pyramidal cells have a smaller specific capacitance, which may to some extent be compensated by the larger size of human neurons (Eyal et al., 2018). Here, we have included distinct human-specific electrophysiological parameters for excitatory and inhibitory cells, but the investigation of further cell-type diversity and the comparison with single-neuron parameters from different species are left to future work.

Various assumptions and approximations flow into the model definition. For instance, with the modeled inhibitory postsynaptic potentials being five times as large as excitatory ones, the relative strength of inhibitory synapses is rather high in the model, in-vitro recordings suggesting a factor closer to 1 (Campagnola et al., 2022). However, reducing the IPSP-to-EPSP ratio even to a value of 2 does not allow adequate reproduction of the observed microscopic and macroscopic activity statistics (see Fig. 9). For simplicity and model robustness, we defined the synaptic strengths via only a few parameters; in reality, synaptic strengths are diverse, for instance having laminar specificity, and the properties of synapses conveying feedforward and feedback signals are likely to differ (Germuska et al., 2006; Bastos et al., 2012; Self et al., 2012). In addition, the electrophysiological properties of individual neurons are known to be distributed, as characterized in detail in the Allen Cell Types Database (Teeter et al., 2018). However, using distributions based on the human neuron parameters provided by the Allen Cell Types Database leads to a worse fit to the experimental data compared to using the mean values only (see Fig. 10). A possible reason is that, in reality, intrinsic neuron parameters and input strengths are attuned to each other, preventing neurons with high intrinsic excitability from being strongly driven (Joseph and Turrigiano, 2017). Another example is that we assumed the fraction of cortico-cortical plus subcortical connections to equal the fraction of white-matter connections; however, cortical areas, especially adjacent ones, may also be connected to some extent via the gray matter (Vandevelde et al., 1996; Anderson and Martin, 2009). Furthermore, the synaptic time constants for excitatory and inhibitory connections are taken to be equal in the model, whereas these have been found to differ in nature (Spruston et al., 1995; Salin and Prince, 1996; Angulo et al., 1999; Gupta et al., 2000). Using longer inhibitory than excitatory time constants does not yield a state where the firing rate and irregularity distributions are simultaneously well reproduced in the present model (see Fig. 11). Further research may show how the model can be made consistent with these various biological features.

Besides qualitative approximations made in the model, detailed parameter values may also be updated in future, as additional data for human cortex are becoming available. For instance, layer- and cell-type-specific connection probabilities, synaptic strengths, and parameters of synaptic dynamics were recently measured in acute slices of human frontotemporal cortex (Campagnola et al., 2022). Furthermore, the recent electron microscopic reconstruction of a millimeter-scale fragment of human temporal cortex (Shapson-Coe et al., 2021) delivers layer- and cell-type-specific local connectivity data that may be used to adjust the microcircuit connectivity used here. The neuron morphologies used here (Mohan et al., 2015, 2023) have important selection effects, being taken from temporal cortex and neurons having to be relatively free of cutting artifacts to be selected for reconstruction, which will tend to favor neurons with relatively small apical dendritic trees. These selection effects may gradually be overcome as new data become available. Enabling further model refinement, a number of valuable resources and results have recently been published, detailing various aspects of histology, immunohistochemistry (Alkemade et al., 2022), transcriptomics (Jorstad et al., 2023; Siletti et al., 2023), and depth-resolved fMRI (Pais-Roldán et al., 2023) of the human brain. Furthermore, detailed human cytoarchitecture and receptor densities are gathered in the BigBrain (Amunts et al., 2013; Wagstyl et al., 2020; Zachlod et al., 2023), and are still being complemented with new measurements. These data follow the Julich-Brain parcellation (Amunts et al., 2020), which is more fine-grained than the Desikan-Killiany parcellation used here. Thus, the data may be leveraged either by finding an appropriate mapping between parcellations or by increasing the granularity of the model.

Experimental functional connectivity is not stationary but exhibits slow fluctuations (Deco et al., 2011). Currently, our model does not exhibit dynamics on such long timescales; we hypothesize that additional slow processes like spike-frequency adaptation, short-term plasticity, or neuromodulation are necessary to this end. Furthermore, the absence of slow activity may lead to an overestimation of the correlations in the functional connectivity estimation when applying the Balloon-Windkessel model or low-pass filtering the signal. To avoid that, we opted to base our fMRI BOLD proxy directly on the summed synaptic inputs. However, it should be noted that a direct comparison of the estimated absolute values with experimental data may not be ideal since we consider shorter timescales in our measure. Therefore, other methods should be explored in the future to account for these issues.

Our model provides a starting point for investigating cortical processes including adaptation, plasticity, and neuromodulation via simulation. It enables in silico studies of the multi-scale dynamics of the human cerebral cortex and the information processing it supports, from the level of spiking neurons to that of interacting cortical areas. Thus, the source code will be publicly available at https://github.com/INM-6/human-multi-scale-model to facilitate rapid model development.

## Acknowledgments

We thank Sebastian Bludau and Timo Dickscheid for helpful discussions about cytoarchitecture and parcellations. Furthermore, we gratefully acknowledge all the shared experimental data that underlies our work, and the effort spent to collect it.

This work was supported by the German Research Foundation (DFG) Priority Program “Computational Connectomics” (SPP 2041; Project 347572269), the European Union’s Horizon 2020 Framework Programme for Research and Innovation under Specific Grant Agreement No. 945539 (Human Brain Project SGA3), the Joint Lab “Supercomputing and Modeling for the Human Brain”, and HiRSE_PS, the Helmholtz Platform for Research Software Engineering - Preparatory Study, an innovation pool project of the Helmholtz Association. The use of the JURECA-DC supercomputer in Jülich was made possible through VSR computation time grant JINB33 (“Brain-Scale Simulations”).

## Appendix

### Mean-Field Theory

While developing the model, it was often beneficial to have a prediction of the activity without performing a computationally demanding full-scale simulation. To this end, we employed the mean-field theory developed by Amit and Brunel (1997) in combination with the extension to exponential postsynaptic currents derived by Fourcaud and Brunel (2002). Within this theory, the input to a neuron in population *A* is approximated as a Gaussian white noise with mean *µ*_*A*_ and noise intensity 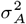. The main assumptions of this theory are that the inputs are uncorrelated, that the temporal structure of the input can be neglected, that a Gaussian approximation of its statistics is valid, that the delays can be neglected, and that the variability of the neuron parameters and synaptic weights can be neglected.

Despite the simplifying assumptions, the theory provides a reliable prediction of the average firing rates if the network is in an asynchronous irregular state (Fig. 8). The remaining deviations are likely a consequence of the neglected variabilities. Thus, the theory allows for rapid prototyping without the need for high-performance computing resources.

**Fig. 8.**
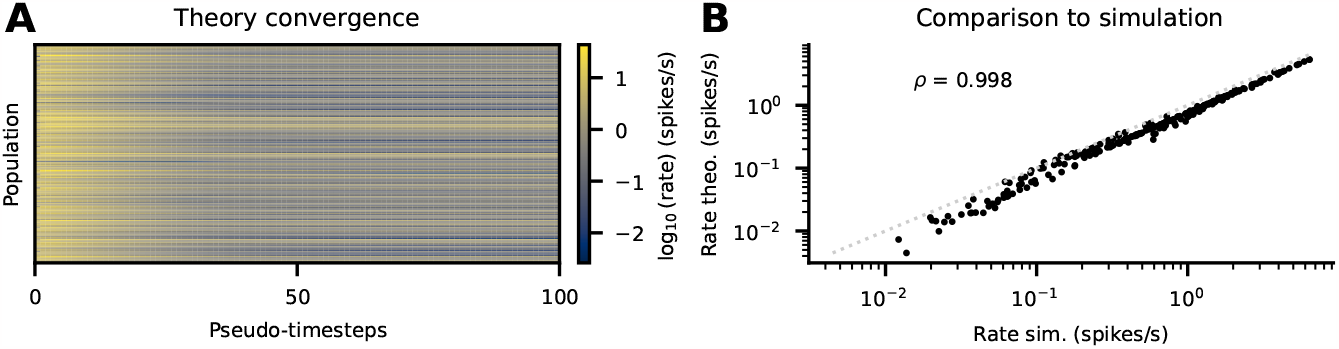
Mean-field theory. (**A**) Firing rates of all populations across the pseudo-timesteps used to find a self-consistent solution. (**B**) Comparison of rates predicted by mean-field theory with empirical rates from a simulation.

## Supplemental Figures

**Fig. 9.**
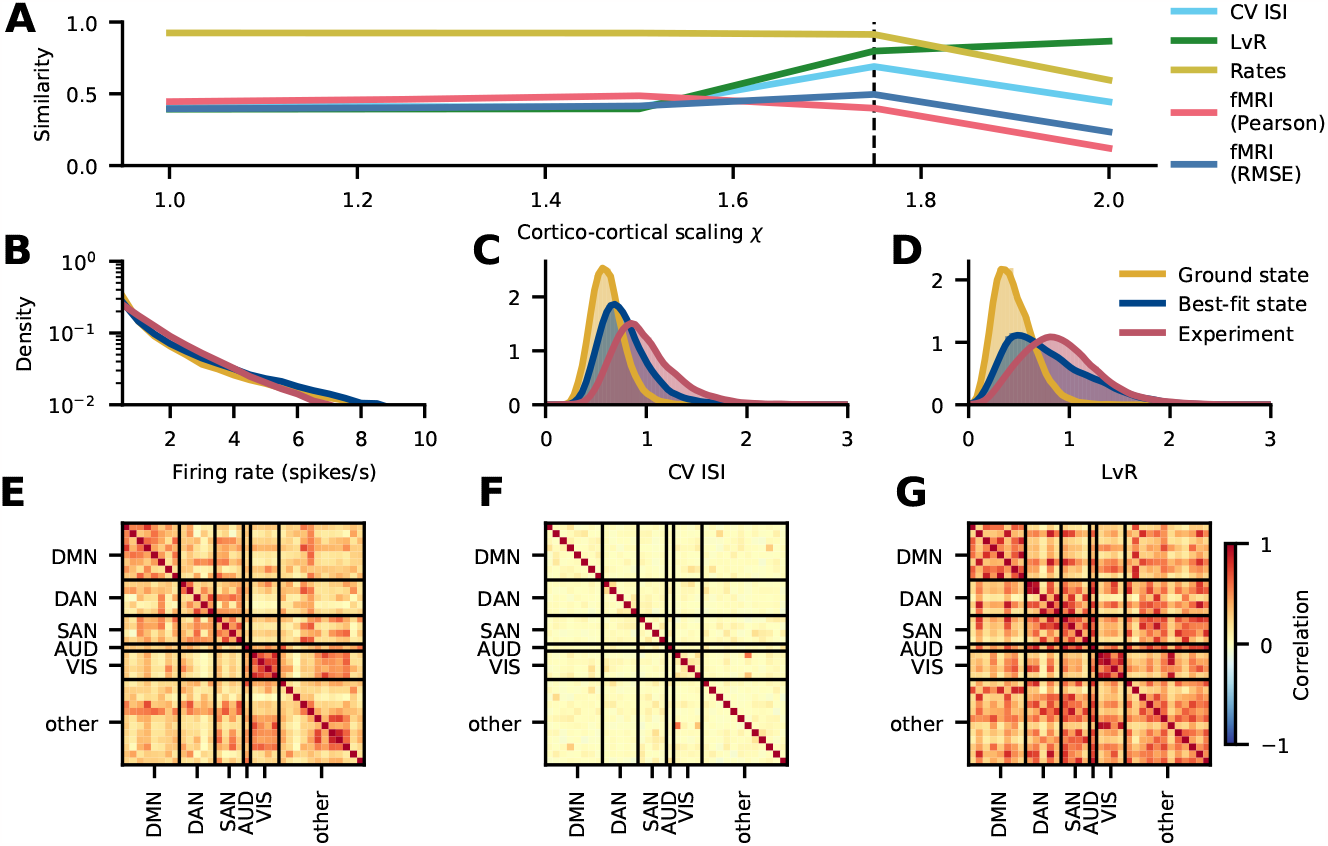
Scaling plot with *g* = *−*2. Simulations abruptly fail at *χ >* 2 and reach a high-activity state that cannot be simulated in a reasonable time. The vertical dashed line at *χ* = 1.75 corresponds to the chosen best-fit state where the agreement with experimental data is maximized. The similarities of the CV ISI, LvR, and rates stay flat across all simulated *χ* while the fMRI similarity slightly increases **(A)**. The firing rate **(B)**, CV ISI **(C)**, and LvR **(D)** distributions of the ground and best-fit states do not differ significantly and, while the rate distributions are close to the experimental data, the irregularity distributions are not. The functional connectivities of the ground state **(F)** and best-fit state **(G)** are weak, unlike the experimental functional connectivity **(E)**.

**Fig. 10.**
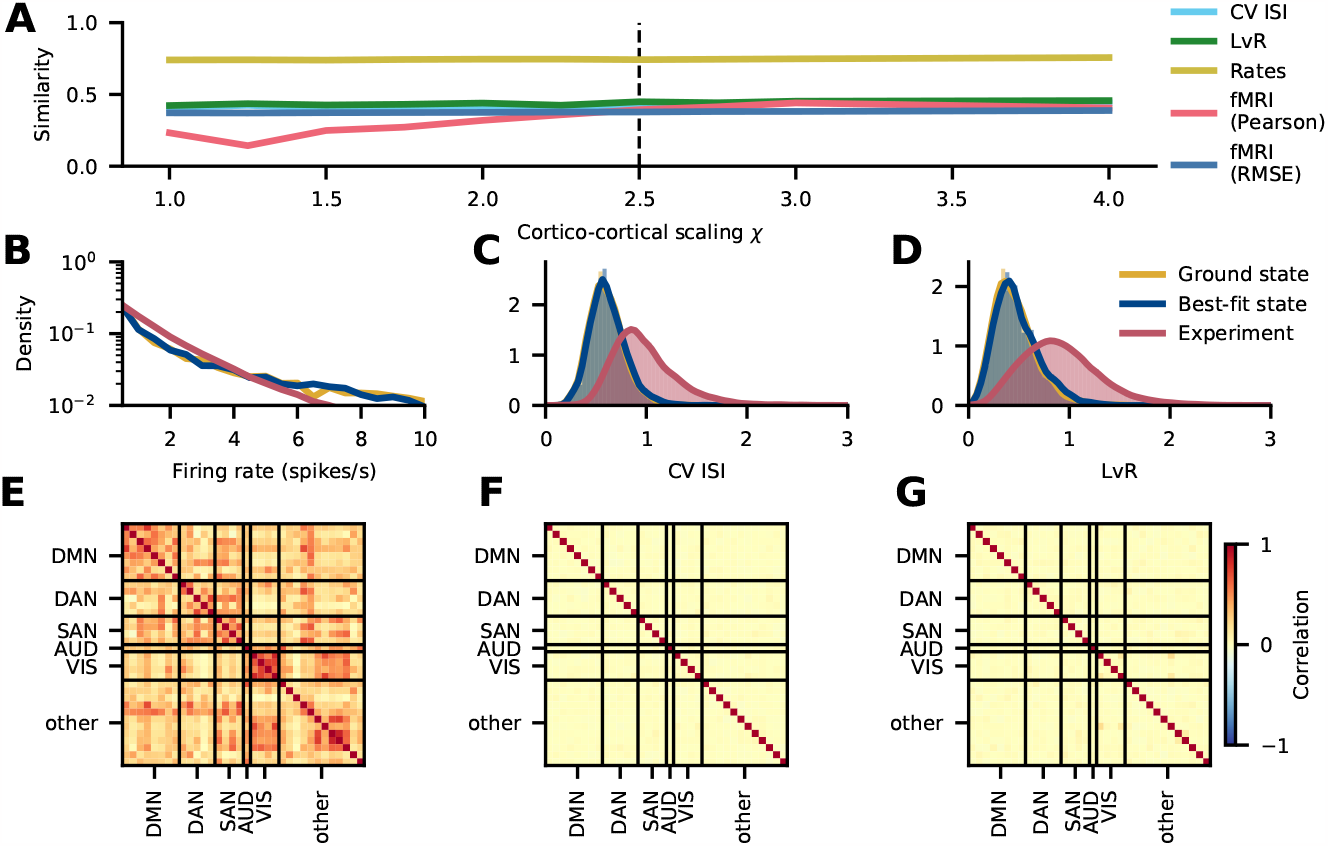
Scaling plot with neuron parameters distributed according to the values for human neurons from the Allen Cell Types Database (https://celltypes.brain-map.org/; cf. Further Model Specifications). As this realization of the model does not show a unique best-fit state, the vertical line at *χ* = 2.5 shows the chosen best-fit state, similar to the original model. While the functional connectivity benefits from increasing the cortico-cortical scaling factors, all other similarities stay constant. Simulations at *χ >* 4 reach a high-activity state that cannot be simulated in a reasonable time. **(A)**. The rates are explained equally well in the ground state and best-fit state, but less well than in the model without distributed neuron parameters (cf. Fig. 5) **(B)**. The CV ISI is not well explained in the ground nor in the best-fit state **(C)**. The LvR in the best-fit state, in contrast to the ground state, matches the experimental data reasonably well **(D)**. The functional connectivity in both the ground state **(F)** and the best-fit state **(G)** is weak, unlike the experimental functional connectivity **(E)**, although the similarity measure indicates that the best-fit state captures the structure of the functional connectivity better.

**Fig. 11.**
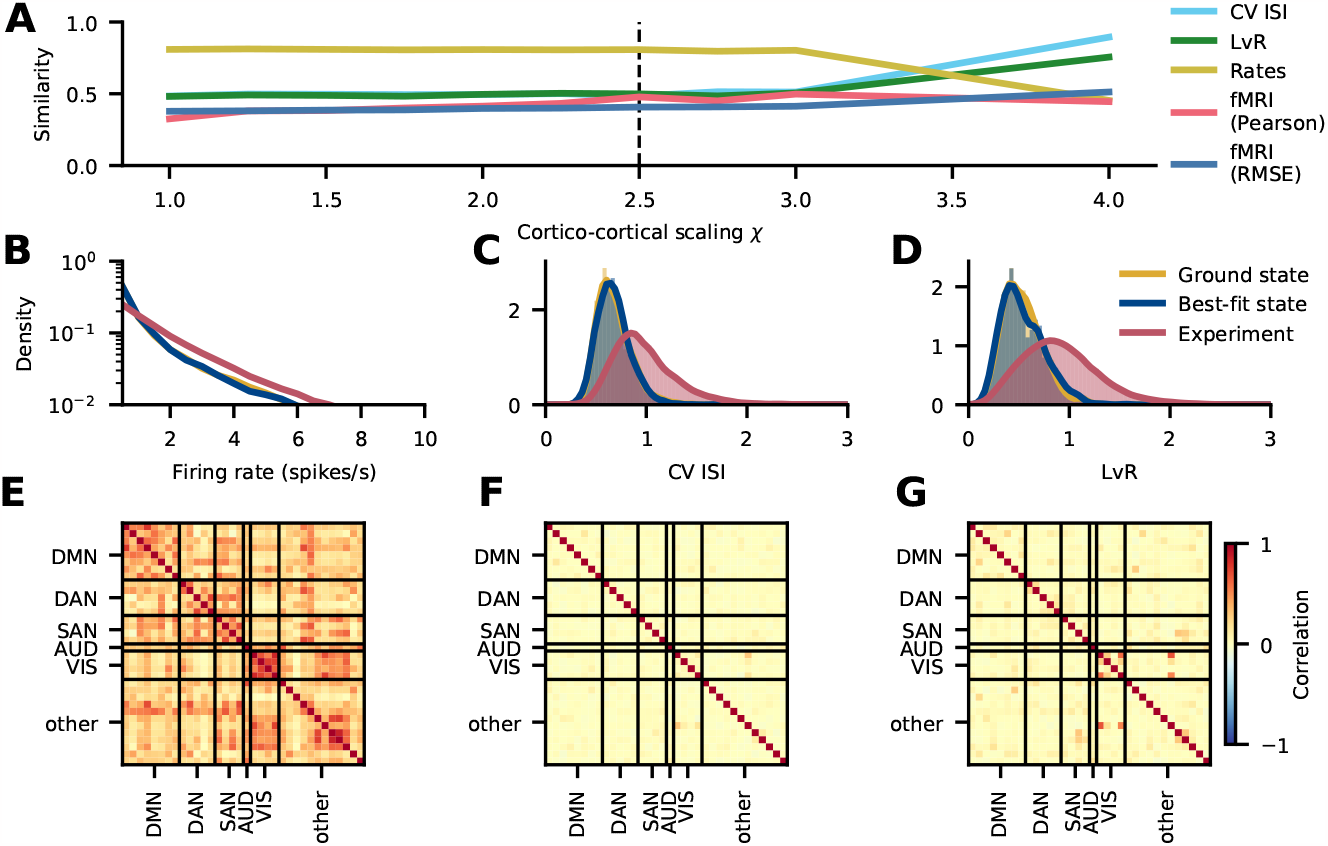
Scaling plot with different synaptic time constants, *τ*_*s,E*_ = 2 and *τ*_*s,I*_ = 4. As this realization of the model does not show a unique best-fit state, the vertical line at *χ* = 2.5 shows the chosen best-fit state, similar to the original model. While the LvR and CV ISI benefit from cortico-cortical scaling factors, the similarity between the simulated and experimental rate distributions degrades and the similarity of the functional connectivity stays constant. **(A)**. The rates are better explained in the ground state **(B)**, whereas the CV ISI **(C)** and LvR **(D)** distributions are nearly the same in both states, and do not match the experimental data. The functional connectivity in both the ground state **(F)** and the best-fit state **(G)** is weak. Neither matches the experimental functional connectivity **(E)** in size, although the similarity measure shows that the structure is fairly well captured.

**Fig. 12.**
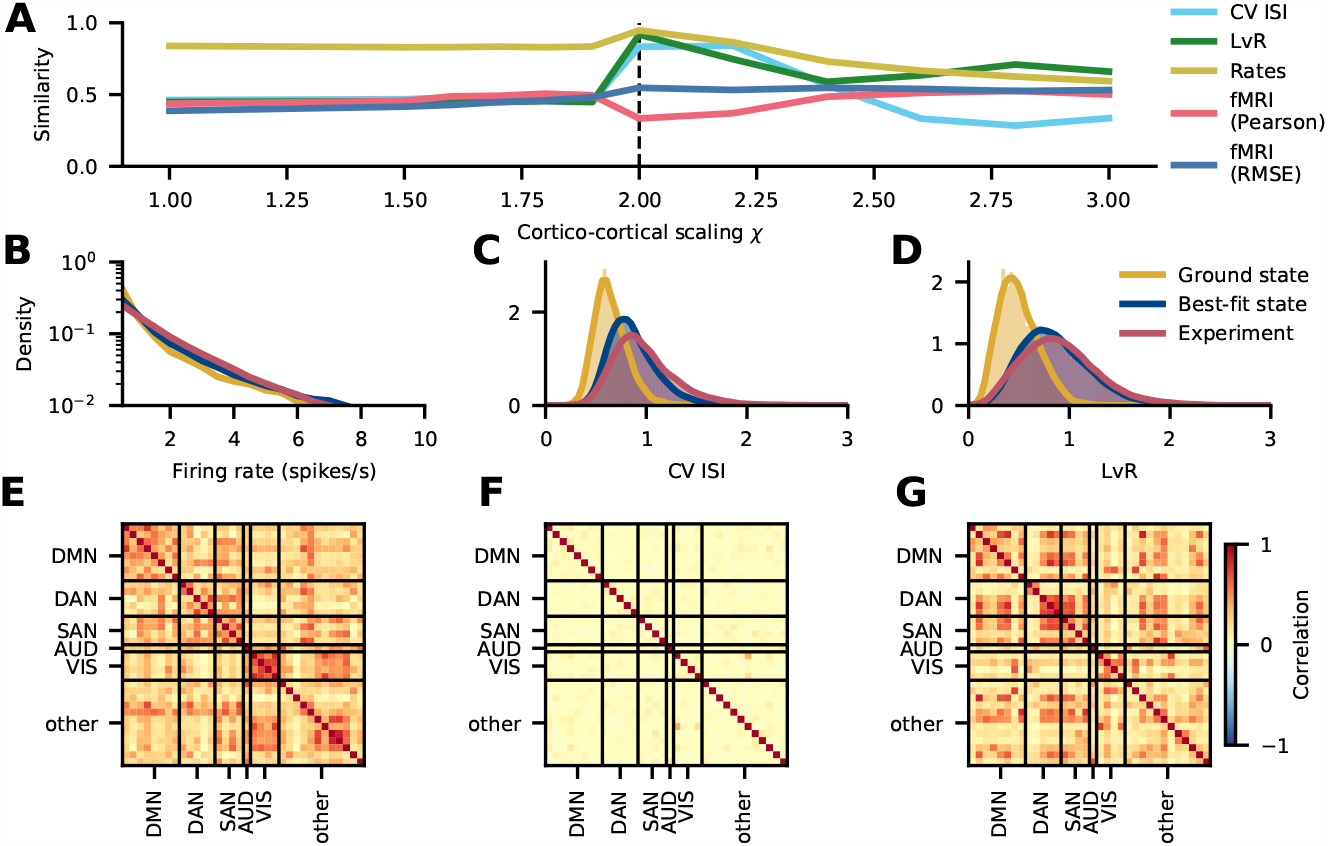
Scaling plot using layer-resolved excitatory-to-inhibitory ratios of neuron densities from Gabbott and Somogyi (1986); Binzegger et al. (2004); Potjans and Diesmann (2014). The main difference compared to Shapson-Coe et al. (2021) is the higher fraction of excitatory cells in layer 2/3: 0.78 vs. 0.65. The other fractions are similar between the two studies: layer 4: 0.80 vs. 0.79, layer 5: 0.82 vs. 0.78, and layer 6: 0.83 vs. 0.86 for Binzegger et al. (2004); Potjans and Diesmann (2014) and Shapson-Coe et al. (2021), respectively. In this model realization, the additional scaling parameter *χ*_*I*_ for cortico-cortical connections onto inhibitory neurons is *χ*_*I*_ = 1. The similarities of rates, CV ISI, LvR, and functional connectivity stay roughly constant for cortico-cortical scaling factors *χ <* 2. The best-fit state is shown as a dashed vertical line at *χ* = 2. Here, the distributions of rates, CV ISI, and LvR almost perfectly match the experimental data, while the similarity of the functional connectivity is slightly worse than in the ground state. For *χ >* 2, the similarities of rates, CV ISI, and LvR slowly deteriorate, while the functional connectivity similarity improves **(A)**. The firing rates match the experimental data well in both states **(B)**, and the CV ISI **(C)** and LvR **(D)** in the best-fit state closely agree with the experimental data. The functional connectivity in the ground state is uniformly weak **(F)**, while it is more differentiated in the best-fit state **(G)**. Interestingly, the pattern of functional connectivity of the ground state matches the experimental functional connectivity **(E)** better than that of the state with the best overall fit.

**Fig. 13.**
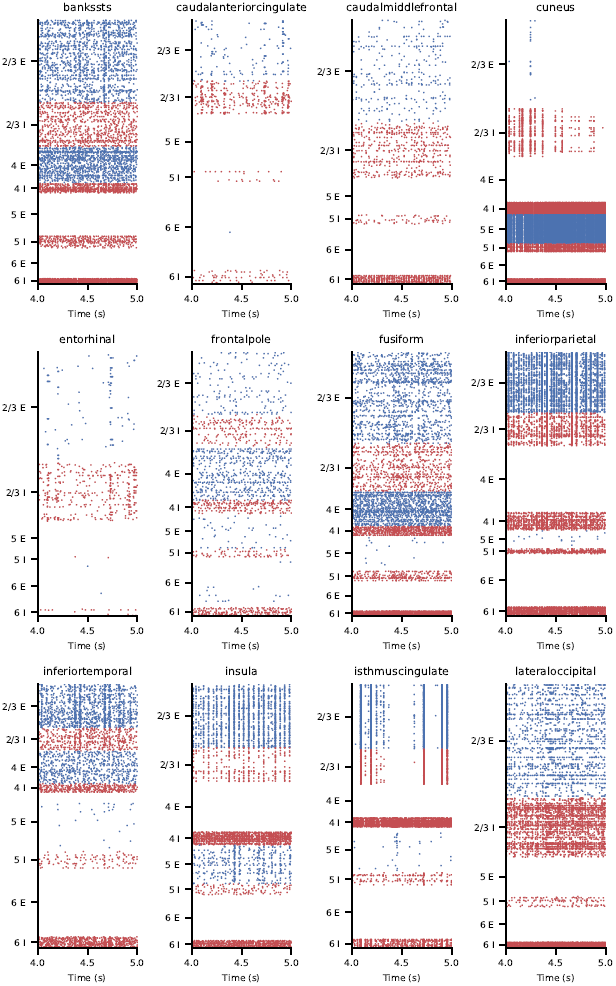
Raster plots in the best-fit state. Subsampled to 2.5% of the excitatory (blue) and inhibitory (red) neurons.

**Fig. 14.**
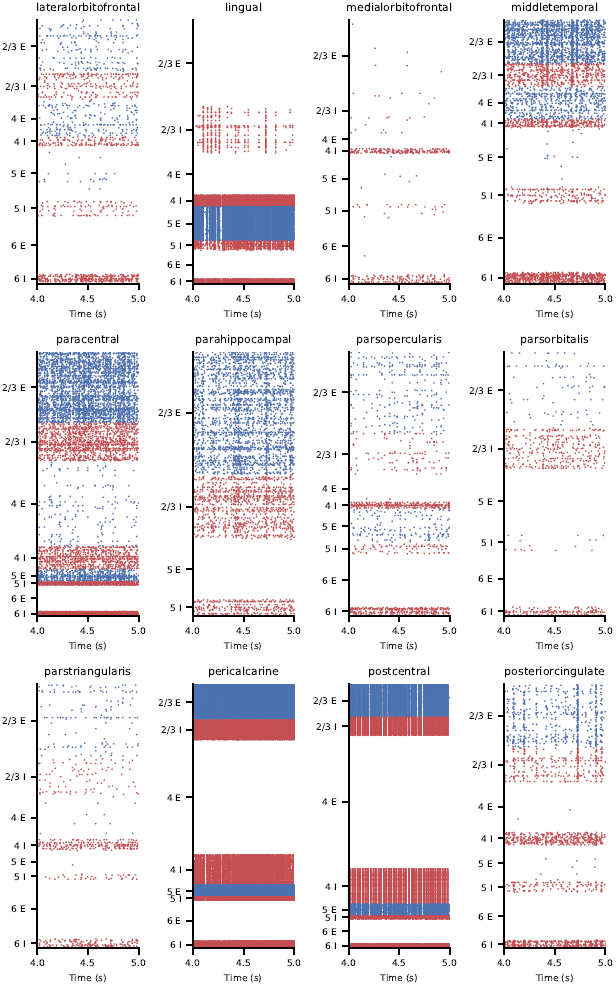
Raster plots in the best-fit state. Subsampled to 2.5% of the excitatory (blue) and inhibitory (red) neurons.

**Fig. 15.**
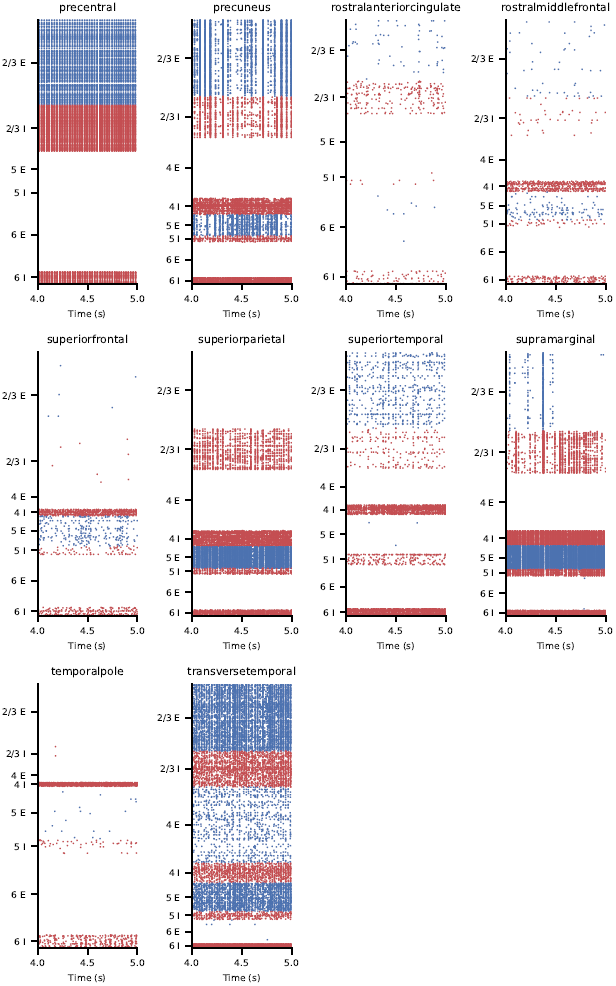
Raster plots in the best-fit state. Subsampled to 2.5% of the excitatory (blue) and inhibitory (red) neurons.

